# Separable codes for read-out of mouse primary visual cortex across attentional states

**DOI:** 10.1101/731398

**Authors:** Ashley M. Wilson, Jeffrey M. Beck, Lindsey L. Glickfeld

## Abstract

Attentional modulation of neuronal activity in sensory cortex could alter perception by enhancing the local representation of attended stimuli or its behavioral read-out downstream. We tested these hypotheses using a task in which mice are cued on interleaved trials to attend visual or auditory targets. Neurons in primary visual cortex (V1) that encode task stimuli have larger visually-evoked responses when attention is directed toward vision. To determine whether the attention-dependent changes in V1 reflect changes in representation or read-out, we decoded task stimuli and choices from population activity. Surprisingly, both visual and auditory choices can be decoded from V1, but decoding takes advantage of unique activity patterns across modalities. Furthermore, decoding of choices, but not stimuli, is impaired when attention is directed toward the opposite modality. The specific effect on choice suggests behavioral improvements with attention are largely due to targeted read-out of the most informative V1 neurons.

## Introduction

In a complex environment with competing incentives, animals must quickly integrate sensory stimuli and flexibly act in a way that depends on current goals. Animals can prioritize specific sensory information through goal-directed selective attention, enabling faster and more sensitive behavioral report of important signals at the expense of less relevant ones (Carrasco, 2011; Maunsell, 2015). Attention is thought to be supported, at least in part, by changes in the neuronal representation of stimuli during sensory processing. Indeed, changes in the firing rate and reliability of responses of visual cortical neurons have been observed during a variety of goal-directed paradigms including spatial (McAdams and Maunsell, 1999; Mitchell et al., 2007; Treue and Maunsell, 1996), feature (Treue and Martinez-Trujillo, 1999; Treue and Maunsell, 1996), and cross-modal attention (Mehta et al., 2000a, 2000b).

Attention-mediated changes in the activity of individual sensory cortical neurons likely contribute to the behavioral effects of attention through their effects on population level cortical computations (Nienborg et al., 2012; Sapountzis and Gregoriou, 2018). Indeed, a major effect of attention is to alter the coordination of population activity (Cohen and Maunsell, 2009; Mitchell et al., 2009), thereby changing how the network represents sensory stimuli across behavioral contexts (Cohen and Newsome, 2008; Lakatos et al., 2009; Raposo et al., 2014; Snyder et al., 2018; Zhang et al., 2011). However, changes in the representation of sensory information may not be sufficient to account for the observed behavioral effects of attention (Krauzlis et al., 2014; Ruff and Cohen, 2018). Instead, contextual changes in population activity may also alter the communication between sensory cortical areas and their downstream targets (Panzeri et al., 2017; Ruff and Cohen, 2016, 2018). Thus, the behavioral effects of attention could be due to changing how efficiently stimulus information is read-out out by downstream areas (e.g. by increasing the efficacy of transmission) in addition to changing the quality of the stimulus information encoded in sensory cortex (e.g. by enhancing the signal-to-noise).

To investigate how attention affects sensory representations and their read-out we monitored populations of neurons in primary visual cortex (V1) of mice while they performed a cross-modal attention task. We find that mice can effectively use a cue at the start of each trial to anticipate either a visual or auditory target. During task performance, V1 neuronal activity is modulated on a trial-by-trial basis such that activity of neurons that encode task stimuli is preferentially enhanced when the mice attend to the visual stimuli. To understand whether changes in sensory responses of V1 neurons could support improved representation or read-out, we decoded the population activity in V1 to predict either the presented stimuli or the animal’s choices. We find that activity in V1 can predict the animal’s choice on both visual and auditory trials, but this prediction is optimized by relying on unique patterns of activity for each modality. Further, the prediction of choice, but not stimulus, is impaired when there is a mismatch between the attended modality and the presented stimulus. The divergence between how V1 represents stimuli and choices across attentional states argues that cross-modal attention modulates which V1 neurons are most effective at driving downstream areas.

## Results

### Mice use a cue to attend to visual or auditory targets in a cross-modal detection task

To understand how sensory cortex supports flexible behavior in a rapidly changing environment we developed a cross-modal detection task for mice. Head-fixed, water-restricted mice were cued on a trial-by-trial basis to expect the appearance of either a visual or auditory target stimulus (Figure 1a-c). Pressing a lever initiated the repeated presentation of a static, vertical (0°), sinusoidal grating (“distractor (D)”; 100 ms duration, 250 ms inter-stimulus interval). The presence or absence of a tone (“cue”; 6 kHz) presented with the first distractor stimulus indicated whether the trial would require the mouse to respond to the visual or auditory target. The presence of the cue indicated that the trial would contain a second target tone (“auditory target (T_A_)”; 10 kHz); conversely, the absence of the cue indicated that the target would be an orientation change (“visual target (T_V_)”). On both trial types, there were at least 2, and up to 10, visual distractor presentations preceding the target. If the mouse released the lever during a short window (100-550 ms) following any target, it was considered a hit and the mouse received a liquid reward.

**Figure 1.**
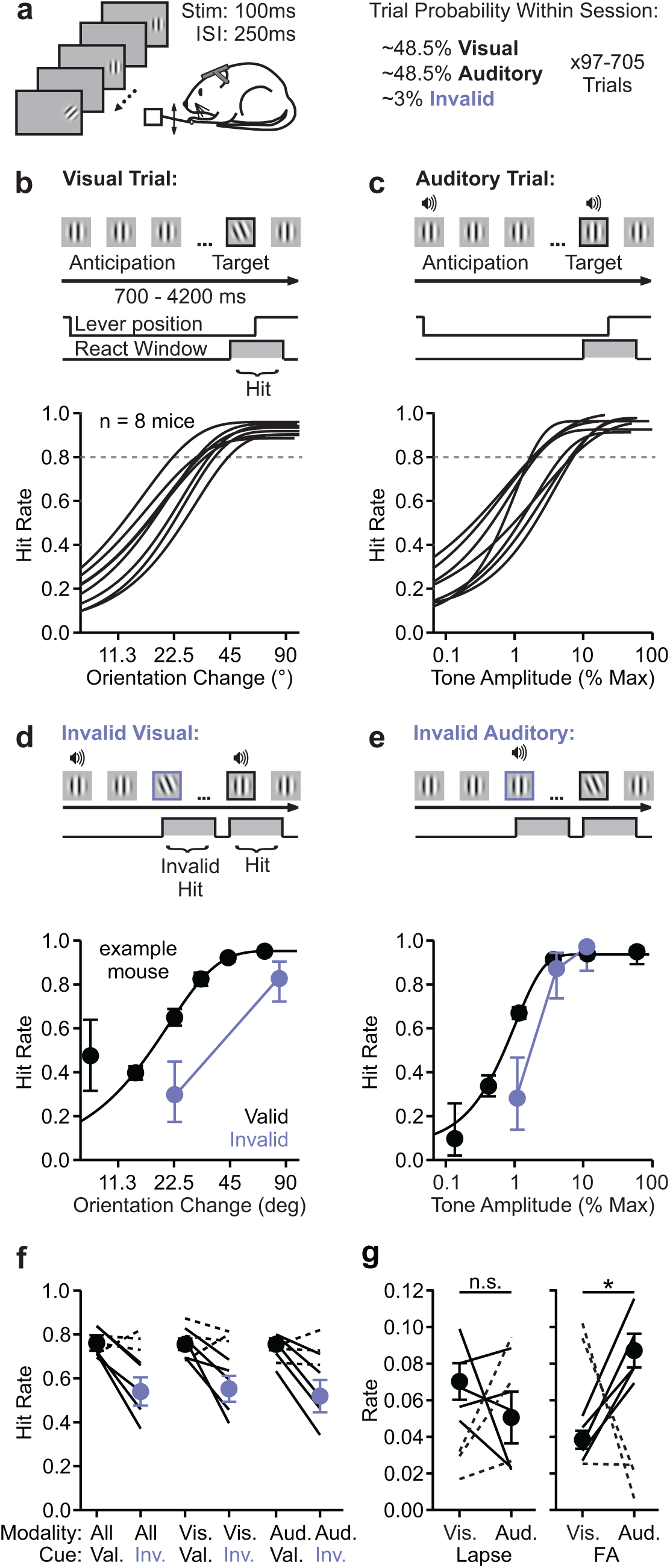
Cues improve detection on a cross-modal change detection task. **a)** Schematic of task structure. Left: Head-restricted mice face an LCD monitor and use a lever to initiate trials and detect targets. Each stimulus (100 ms) is separated by a 250 ms inter-stimulus interval (ISI). Right: Validly-cued visual and auditory trials are presented with equal probability along with very rare invalidly-cued trials. **b)** Top: Schematic of visual trial structure. Upon trial initiation, 2-10 vertical distractor gratings (“Anticipation”) and an orientation change (“Target,” between 8 and 90 degrees) are presented. Mice must hold the lever through the distractors and release the lever within a brief window following target to receive a reward. Bottom: Weibull functions fit to the hit rate on visual trials as a function of target orientation for all mice trained on this task (n = 8). **c)** Same as **b** for auditory trials. Trial structure is the same, except the first distractor is accompanied by a tone (“cue”, 6 kHz), and the target is a second tone (10 kHz). Weibull functions are fit to the hit rate on auditory trials as a function of tone amplitude. **d)** Top: Schematic of an invalidly-cued visual trial. The trial is cued as auditory (tone accompanies first distractor), but a visual target is presented. Lever releases in response to either invalidly- or validly-cued targets are rewarded (see Methods). Bottom: Hit rate across valid (black) and invalid (purple) visual targets for an example mouse. Target orientations are binned by difficulty. Hit rate error is 95% C.I.; target orientation error is S.E.M. **e)** Same as **d** for invalidly cued auditory trials. **f)** Hit rate on valid and invalid trials for each mouse, and average across mice with a significant difference (n = 5). Hit rate was calculated across valid and invalid trials matched for difficulty across all (left), visual (middle), or auditory (right) trials. Solid lines indicate that hit rate was significantly different across all valid and invalid trials (p < 0.05, binomial test). Error is S.E.M. across mice. **g)** Lapse rate (left; p=0.28, paired t-test) and false alarm rates (p<0.001) on visual and auditory trials. Only mice with a significant effect of attention cue included in average. Error is S.E.M. across mice.

We controlled task difficulty by varying the target orientation (8-90°) on visual trials or the amplitude of the target tone (0.03-100% of maximum amplitude) on auditory trials (Figure 1b-c). Probing the animals’ detection thresholds made the task challenging and incentivized the mice to use the cue to attend to the expected target modality. To test whether the mice used the cue to guide their behavior, we presented rare (2.4±0.13% of trials), invalidly-cued visual or auditory trials in which the cue incorrectly predicted the target modality (Figure 1d-e). For instance, on an invalidly-cued visual trial, a tone accompanied the first stimulus presentation indicating that the mouse should expect an auditory target, however a visual target was presented (Figure 1d). On these trials, lever releases within the reaction window following invalidly-cued targets were rewarded; however, if the mouse failed to detect the invalidly-cued target, the trial was allowed to continue and the mouse had the opportunity to detect a valid target.

We compared hit rates for validly and invalidly-cued targets to determine whether the cue improved target detection of the expected modality. Across sessions, five out of eight mice had a significantly lower hit rate on invalidly-cued trials compared to validly-cued trials of comparable difficulty (p<0.0001; binomial test; solid lines Figure 1d-f). These effects were also consistent within sessions (mice with attention all p<0.001, paired t-test; Supplemental Figure 1). Moreover, the five mice with a lower hit rate on invalidly-cued trials had a lower hit rate for invalidly-cued trials of both modalities, consistent with bidirectional effects of the cue on behavior (visual: 4/5 mice p<0.05 (fifth mouse has trend, p= 0.12); auditory: 5/5 mice p<0.05; binomial test; Figure 1g). We considered only these five animals for further analyses.

Factors other than expected target modality, such as sensory input, arousal, reward expectation, and motor planning, should be the same on auditory and visual trials during the interval between the cue and target presentation (“anticipation” phase). Indeed lapse rate (defined as 1 - hit rate for the easiest condition of each modality; visual: 0.076±0.011; auditory: 0.042±0.009; n = 5 mice; Figure 1g) and false alarm rate (FA: lever releases that occurred within the same reaction window as used for hits, but following a distractor; visual: 0.039±0.003; auditory: 0.083±0.009; Figure 1g) were low suggesting consistent levels of task engagement within sessions and across trial types. As a separate measure of task arousal and sensory input, we monitored pupil size and position as the mice performed the task (Supplemental Figure 2a; see Methods). There were slow changes in pupil size during the anticipation phase of the trial, however, there were no consistent differences between visual and auditory trials (p=0.80; paired t-test; Supplemental Figure 2b-c). Similarly, while there were eye movements at the start of the trial and at the time of the target, the deviation (range: 0.05-2.0°) was much smaller than the size of a V1 receptive field (Bonin et al., 2011), and not consistently different between visual and auditory trials (anticipation: p=0.94; target: p=0.47; paired t-test; Supplemental Figure 2c-g). Thus, the cue and the subsequent shift of attention of the target modality are the major factors that differ during the anticipation phase across visual and auditory trials.

### V1 neurons are driven more strongly during the anticipation phase on visual trials

To investigate how attention modulates activity in sensory cortex we used two-photon calcium imaging to monitor the activity of layer 2/3 neurons in V1 virally expressing GCaMP6m as the mice performed the cross-modal detection task (Figure 2a). Cells that responded to visual stimuli were selected from each recording session in post-hoc analyses (n=1367 cells, 5 mice, 14 sessions; see Methods, Figure 2a-c). We focused our analyses on the anticipation phase of the trial, when sensory input and behavioral state are similar across trial types, but attention is directed toward visual or auditory stimuli. The dynamics of neuronal responses to the repeated distractor presentations were diverse (Figure 2b-c). Many neurons were significantly driven by the first distractor stimulus (n=418/1367 neurons; Figure 2b-c; example neuron 1), whereas others were only significantly driven late in the trial (n=245/1367 neurons; Figure 2b-c; example neuron 2) or only suppressed late in the trial (n=291/1367 neurons, Figure 2b-c; example neuron 3).

**Figure 2.**
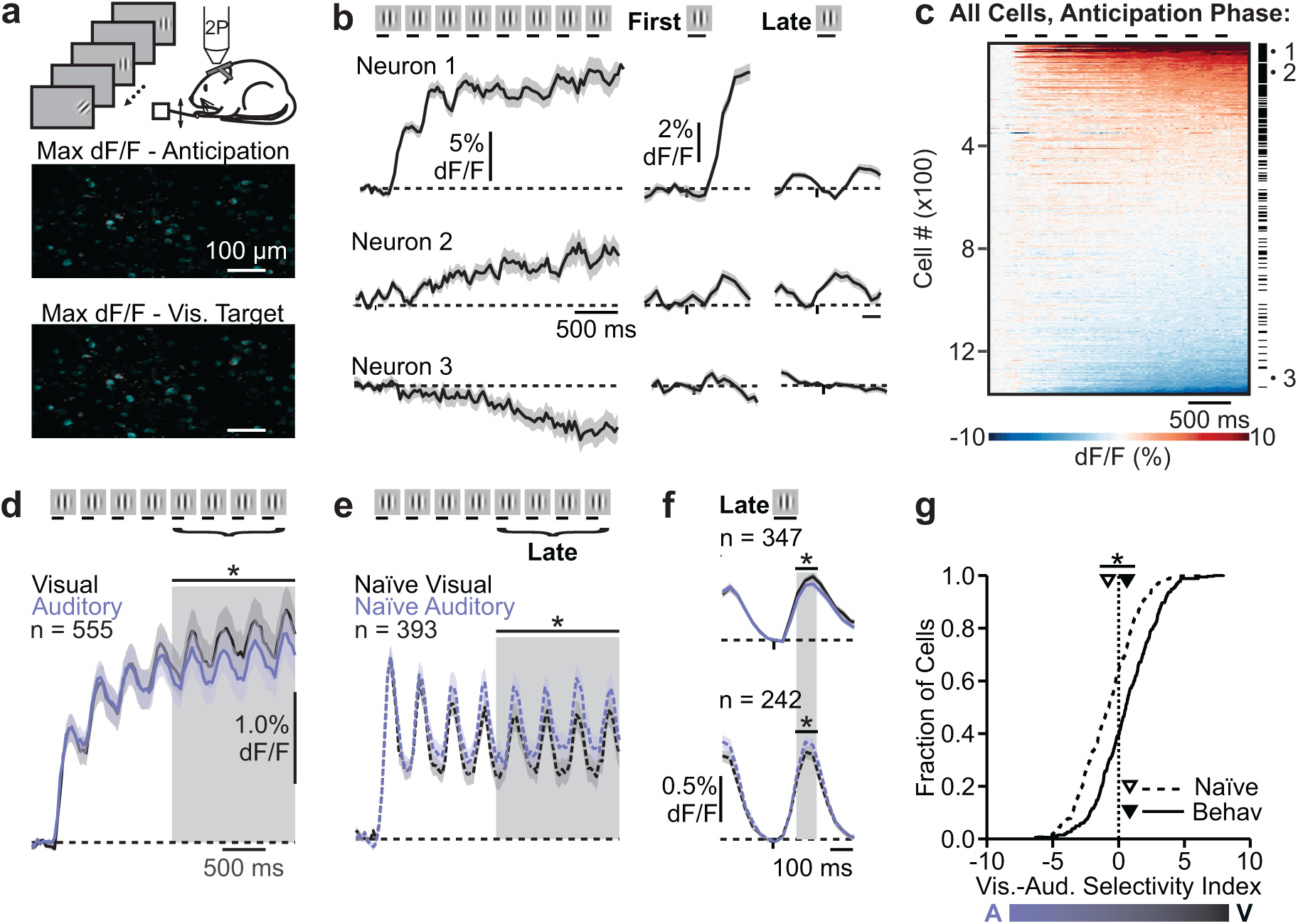
V1 neurons are modulated by attentional state. **a)** Top: mouse performs the attention task during two-photon (2P) imaging. Middle: field of view image of maximum change in fluorescence (max dF/F) during the anticipation phase of the trial. Bottom: max dF/F image from target phase of the trial. Overlay of selected cell masks in cyan. **b)** Left: Time-course of activity during the anticipation phase for all trials with at least 8 distractors (left), all first distractor (middle), or all late distractor stimuli (5^th^-10^th^, right) for three example V1 neurons. Shaded error is S.E.M. across trials. **c)** Heatmap of dF/F during the anticipation period for trials with at least 8 distractors for all neurons (n = 1367, 5 mice) recorded during behavior sorted by response amplitude during the late anticipation period. Tick marks indicate anticipation responsive cells; #1-3 are the cells in **b**. **d)** Average time-course of anticipation responsive neurons on visual (black) and auditory (purple) trials with at least 8 distractors. Only neurons from experiments with 100 ms stimulus duration are shown (1096/1367, see Methods). Shaded gray area is analysis window. Significance is tested across all neurons (663/1367); *p<0.0001, paired t-test. Error is S.E.M. across cells. **e)** Same as **d**, for a separate cohort of naïve mice (n=393/633, 9 mice); *p<0.001, paired t-test. **f)** Top: average time-course of all late anticipation responsive neurons to all late distractor stimuli for behaving mice (n=347/1367). Shaded gray area is analysis window. Bottom: Same as top, for naive mice (n=376/633). *p<0.001, paired t-test. **g)** Cumulative distribution of visual minus auditory selectivity index (SI_VA_) of responsive neurons from behaving or naïve mice calculated from late distractor stimuli shown in **f**. Arrows indicate mean SI_VA_ for behaving and naïve mice. *p<0.0001, Kolmogorov-Smirnov test.

Because responses on auditory trials reflect the population activity that accompanies impaired detection on invalidly cued visual trials we compared neuronal activity on visual and auditory trials to determine how attention impacts visual responses in V1. Indeed, we found a reliable increase in visually-driven activity in V1 neurons when the visual stimulus was attended (Figure 2d, f). On average, we find that anticipation-responsive V1 neurons (i.e. responsive to the first stimulus or late in the trial) had greater responses on visual trials as compared to auditory trials, but only late in the trial (early window (0-1400 ms): p=0.54; late window (1400-2833 ms): p < 0.0001; paired t-test, n=663 neurons; Figure 2d). Differences across trial types during this late phase incorporate both time-locked, visually-driven responses as well as slower, sustained changes in activation. To quantify how visually-driven responses to the distractor change with attention, we aligned the onset of each distractor stimulus occurring late in the trial (fifth through last distractor before the target) and identified the subset of responsive cells that were reliably driven by these late distractors (“late-responsive”; n = 347 cells; Figure 2f). Late-responsive neurons had greater visually-driven responses on attend-vision trials (p<0.001, paired t-test; Figure 2f, top) and a mean visual-auditory selectivity index (SI_VA_; the difference between each neuron’s variance normalized average response on visual trials and auditory trials) that was significantly greater than zero (p<0.0001, Student’s t-test; Figure 2g). Thus, visually driven neuronal responses in V1 are enhanced when the mouse is attending to visual targets.

Differences in neuronal activity across trial modality cannot be explained by sensory effects of the auditory cue. To investigate the sensory contribution of the cue in the absence of attention, we imaged V1 neurons in a separate cohort of naïve mice as they passively viewed a movie of the cross-modal task. In these naïve mice, distractor-driven V1 neuron responses were actually smaller on visual trials compared to auditory trials (n=393 neurons, p<0.001, paired t-test; Figure 2e). Similarly, in these mice, visually-driven responses were larger on auditory trials (n=242 neurons, p<0.0001, paired t-test; Figure 2f, bottom), and across neurons the average SI_VA_ was less than zero (p<0.0001, Student’s t-test; Figure 2g). Moreover, while there were similar proportions of significantly modulated cells in behaving and naïve mice, we found that there were many more +SI_VA_ neurons in the behaving mice (behavior – 15.0% modulated, 38/52 with +SI_VA_ I; naïve – 16.1% modulated, 5/39 with +SI_VA_, Supplemental Figure 3a). Thus, observed increases in V1 neuron activity during the anticipation phase of the task are due to changes in selective attention, and may even be competing against the suppressive effects of multi-sensory interactions.

Finally, we addressed the population of cells that were suppressed during the late phase of the anticipation period. In this population, we found no significant difference between attend-visual and attend-auditory trials during the late response window (dF/F visual: -2.3±0.1%; dF/F auditory: -2.2±0.1%; p=0.37, paired t-test, n=291 cells; Supplemental Figure 4). There were neurons that were significantly modulated across attentional conditions (11.1% modulated), however there was a similar proportion of modulated suppressed cells in the naïve dataset (10.9% modulated) and the fraction of +SI_VA_ and -SI_VA_ cells were balanced within both groups (behaving: 16/35 with +SI_VA_; naïve: 2/5 with +SI_VA_, Supplemental Figure 3a). Thus, we conclude that attention increases the activity of driven V1 neurons and has no net effect on the activity of suppressed cells.

### Visual attention increases activity in task-relevant neurons

While distractor-responsive V1 neurons had a greater average response on visual trials, there was substantial diversity in the magnitude and direction of attentional modulation. Thus, we next sought to determine whether neurons’ functional properties could explain this diversity in modulation. We hypothesized that attention may preferentially increase responses of V1 neurons that are useful for performing the task. Because orientation tuning is an important feature for differentiating targets and distractors, we next analyzed responses to drifting gratings presented in a passive session following the behavior. These experiments allowed us to measure each neuron’s full orientation tuning curve (0-180°) because task orientations were only varied between 0-90° (Figure 3a). We then binned late-responsive neurons by their preferred orientation (0°, n=41; 45°, n = 60; 90°, n = 81; 135°, n = 67) to test the hypothesis that attentional modulation is specific to task-informative neurons.

**Figure 3.**
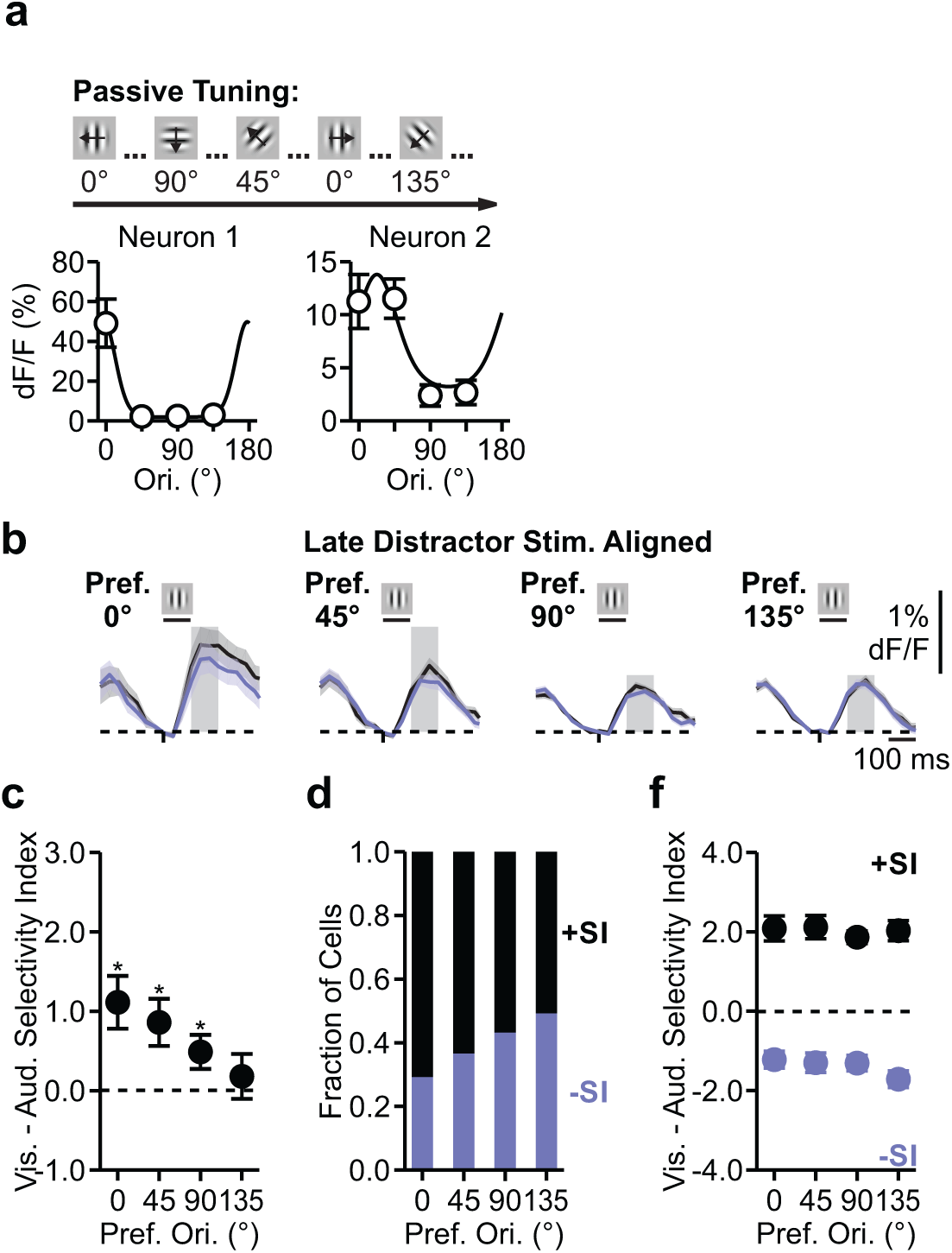
Attention modulation depends on orientation tuning. **a)** Schematic of passive orientation tuning experiment and orientation tuning of two example neurons. Curves are von Mises fits. Error is S.E.M. across trials. **b)** Average time-course of late distractor response to for the same groups of neurons in. Error is S.E.M. across cells. **c)** Mean SI_VA_ for each group shown in **c**. *p<0.05, Student’s t-test. Error is S.E.M. across cells. **d)** Fraction of cells in each orientation preference group with positive or negative SI_VA_. **e)** Mean positive or negative SI_VA_ for each orientation preference group.

Neurons’ attentional modulation depended on their orientation preference. On average, only neurons with orientation preference that matched the task stimuli (i.e. 0°-, 45°-, and 90°-preferring neurons) had a significantly positive attentional modulation (average SI_VA_, 0°: p<0.01; 45°: p<0.01; 90°: p<0.05; 135°: p=0.53; Student’s t-test, Figure 3b-c). This difference in the magnitude of attention modulation was largely explained by the fraction of neurons with positive or negative selectivity within each group. Zero-preferring neurons had significantly more positively than negatively modulated neurons and 45°- and 90°-preferring neurons showed the same trend (0°: p<0.05; 45°: p=0.14; 90°: p=34; 135°: p=0.86, Chi-squared test; Figure 3d). The magnitude of selectivity was similar across orientation preference groups when positive and negative selectivity neurons were assessed independently (positive-p=0.36; negative-p=0.87; one-way ANOVA; Figure 3e). Thus, neurons that prefer task orientations are more likely to be positively modulated by attention than those that prefer orientations not used in the task.

### Information about visual and auditory stimuli and choices is present in V1

Two major (non-mutually exclusive) hypotheses for the neuronal basis of attention posit that its behavioral effects could be due to enhancement in 1) the signal-to-noise of the sensory cortex population response or 2) the efficiency of the downstream decoder in reading out that activity. The observed attentional modulation of the activity of task-relevant V1 neurons might support either hypothesis, either through direct impact on the stimulus representation or read-out of V1, or by reflecting changes in the downstream decoder. Thus, we took a regression-based approach to test how attention-dependent changes in V1 neuronal activity alter representation and read-out of sensory stimuli. Specifically, we assessed the accuracy of representation in V1 by decoding the activity of simultaneously recorded populations of neurons in response to single stimulus presentations during behavior to predict the whether the stimulus was a target or distractor (“stimulus model,” Figure 4a-b). We also used the same neuronal population responses to predict whether the choice was yes or no (“choice model,” Figure 4a, c) as a measure of the accuracy of read-out. Differences in the performance of these models across visual and auditory trials reflect differences in how task information can be extracted from V1 neurons and can thus reveal how modulation of V1 neuron populations might support changes in attention across modalities.

**Figure 4.**
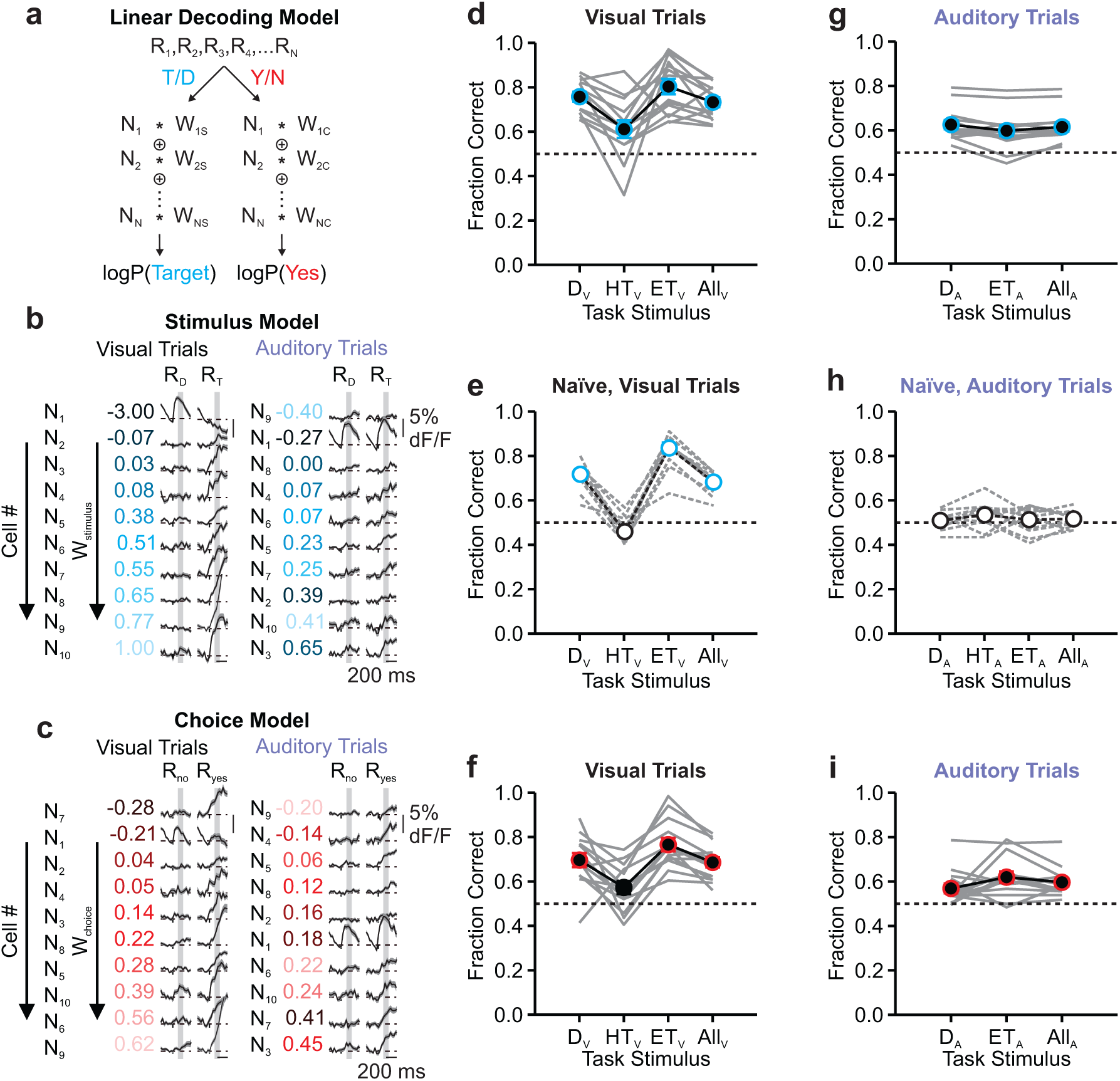
Stimuli and choices on both visual and auditory trials can be predicted from V1 population activity. **a)** Linear models were trained to discriminate between targets (T) and distractors (D) (“Stimulus Model”, blue) or yeses (Y) and noes (N) (“Choice Model”, red) using responses (R_1_-R_n_) from populations of neurons (N_1_-N_n_), assigning a weight (W_1_-W_n_) to each for predicting which type of stimulus or choice occurred (logP). **b)** Average response of modeled neurons from an example session to visual (left) or auditory (right) distractors (R_D_) and targets (R_T_), sorted by weight in the visual stimulus model. Neurons are color-coded and have the same number ID (N_n_) across visual and auditory stimulus models. Shaded error is S.E.M. across trials. **c)** Same as **b** for responses to noes (R_no_) or yeses (R_yes_), sorted by weight in the visual choice model. Neurons have the same number ID (N_n_) across stimulus and choice models. **d)** Fraction of stimuli correctly identified when trained and tested on visual trials, binned by stimulus type (distractors (D_V_), Hard Targets (8-32°, HT_V_), and Easy Targets (33-90°, ET_V_)) or combined across all trials. Solid gray lines connect data from individual imaging sessions (n=14). Colored points indicate that the model performed significantly better than chance (0.5) across experiments; all points: p<0.05. Error is S.E.M. across experiments. **e)** Performance of the visual stimulus model when trained with neurons from naïve mice (n=11 sessions). Colored points: p<0.0001. **f)** Same as **d**, for the predicting the animal’s choice on visual trials. Colored points: p<0.0001. **g-i)** Same as **c-f** for stimulus (all points: p<0.001), naïve stimulus, and choice (all points: p<0.001) models trained and tested auditory trials. Note that HT_A_ were not tested due to low numbers in some experiments.

For both models, we selected a subpopulation of strongly task-driven (i.e. target- or distractor-responsive), orientation tuned neurons from each imaging session, trained a generalized linear model to discriminate the stimulus or choice for all but one presentation, and then used the fit weights to test held-out presentations (see Methods). The population of V1 neurons performed well above chance at predicting the type of visual trial stimulus (distractor (D) or target (T), All_V_-p<0.0001; Student’s t-test; n = 5 mice, 14 sessions; Figure 4d). The model performed above chance for all visual stimulus types (Distractor (D_V_): p<0.0001; Hard Target (HT_V_): p<0.05; Easy Target (ET_V_): p<0.0001; Student’s t-test) but significantly worse on hard visual targets (one-way ANOVA (p<0.001) with post-hoc Tukey test: hard target compared to all others: p<0.01). In naïve mice (untrained animals, passively viewing task stimuli), the stimulus model performed well above chance across all presentations (p<0.0001; Student’s t-test; Figure 4d), but performed at chance at detecting hard visual targets (D_V_: p<0.0001; HT_V_: p=0.98; ET_V_: p<0.0001; Student’s t-test). Thus, there is information in V1 about whether the stimulus is a visual target or distractor. Further, information about the stimulus is enhanced in the behaving condition.

We also found that there is information in V1 that can be used to predict the mouse’s choice on visual trials: whether it responded yes (i.e. hits and false alarms) or no (i.e. misses and correct rejects; All_V_-p< 0.0001; Student’s t-test, Figure 4c, f). Similar to the stimulus model, the choice model performed well on visual distractor (p<0.0001) and easy visual target (p<0.0001) test stimuli, but performed worse on hard visual targets (p=0.029 compared to chance, Student’s t-test; one-way ANOVA (p<0.01) with post-hoc Tukey test: hard target compared to others: p<0.05). This demonstrates that fluctuations in V1 neuron activity are tightly linked to perception of the visual stimulus.

To address how these population representations might change across attentional conditions, we next trained a model to discriminate the auditory distractor (D_A_) from target (T_A_) stimuli. Notably, since the visual stimuli accompanying auditory distractors and targets are identical (i.e. vertical gratings), and V1 neurons are not known to explicitly represent auditory stimuli, we did not expect that there would be information in V1 for discriminating auditory targets from distractors. Thus, it was surprising that we were still able to discriminate auditory targets and distractors above chance on auditory trials (All_A_-p<0.0001, Student’s t-test; Figure 4g), and the model performed only slightly better at predicting auditory targets than distractors (paired t-test, p<0.05). Unlike the visual condition, data from naïve animals could not be used identify auditory stimuli (p=0.18; Student’s t-test; Figure 4h). This suggests that behaving or training in this task gates the propagation of information about the auditory stimulus into V1.

As with visual trials, we also found that there is information in V1 about the mouse’s choices on auditory trials (All_A_: p<0.0001; D_A_: p<0.01; ET_A_: p<0.001; Student’s t-test; Figure 4i). The ability to predict choice cannot simply be explained by signals in V1 reflecting whether the mouse was rewarded or not, since many of the presentations in which the mouse responded “yes” were not rewarded (i.e. correct rejects). However, a trivial explanation for these signals could be a motor feedback signal, since all “yes” responses involve releasing the lever. To address this possibility, we analyzed the performance for each stimulus and choice model in varying time windows following stimulus presentation. For both the visual and auditory trials, both stimulus and choice information could be predicted before the earliest allowed reaction time (minimum reaction time: visual = 200ms, auditory=150ms; time when model performance is above 55% correct: 52±6ms, visual detect: 60±10ms, auditory target: 98±14ms, auditory detect: 110±15ms; n=14 sessions; Supplemental Figure 4). Thus, neuronal activity in V1 is sufficient to predict both the stimulus presented and the animal’s choice on both visual and auditory trials. Moreover, the model is likely using sensory signals, and other signals that precede the decision, rather than motor or reward-related activity to discriminate choice.

### Linearly separable codes for visual and auditory stimuli and choices in V1

The presence of a population code in V1 for auditory stimuli and choices is only surprising if it is truly independent from the population code for visual stimuli and choices. While our analysis argues against a contribution of reward or motor-related signals, there are other shared sensory signals that might contribute to the prediction of auditory information from V1 activity. For instance, efferent copy or reward prediction signals might significantly precede the movement. In addition, visual and auditory distractors are identical (i.e. both are vertical gratings without an auditory tone), and therefore information supporting the identification of visual distractors might contribute to the identification of auditory distractors. If the discrimination of auditory stimuli and choices were due to such shared signals, then we would expect a strong correlation between the weights of each neuron for predicting visual or auditory information. However, auditory and visual weights were only weakly correlated with each other when predicting either the stimulus or the choice (stimulus: explains 0.95% of the variance; R=0.097, p<0.05; Figure 5a; choice: explains 4.8% of variance; R=0.22, p<0.0001; Pearson correlation; Figure 5b). To further measure the independence of weights across modalities, we measured the vector angle across visual and auditory weights for both stimulus and choice models. If the weights were significantly correlated, then we would expect a peak at π/4 (where auditory and visual weights are equal). However, the distribution of vector weights was relatively flat, suggesting that weights are independent across modalities (Figure 5c-d). One possible explanation for this weak correlation would be if there are many possible solutions to the regression. However, this is not the case since we find that the weights for stimuli and choices were both highly correlated within modality (visual-R=0.63; auditory-R=0.80; Supplemental Figure 5a-b). This suggests that there are separable codes in V1 for visual and auditory stimuli and choices.

**Figure 5.**
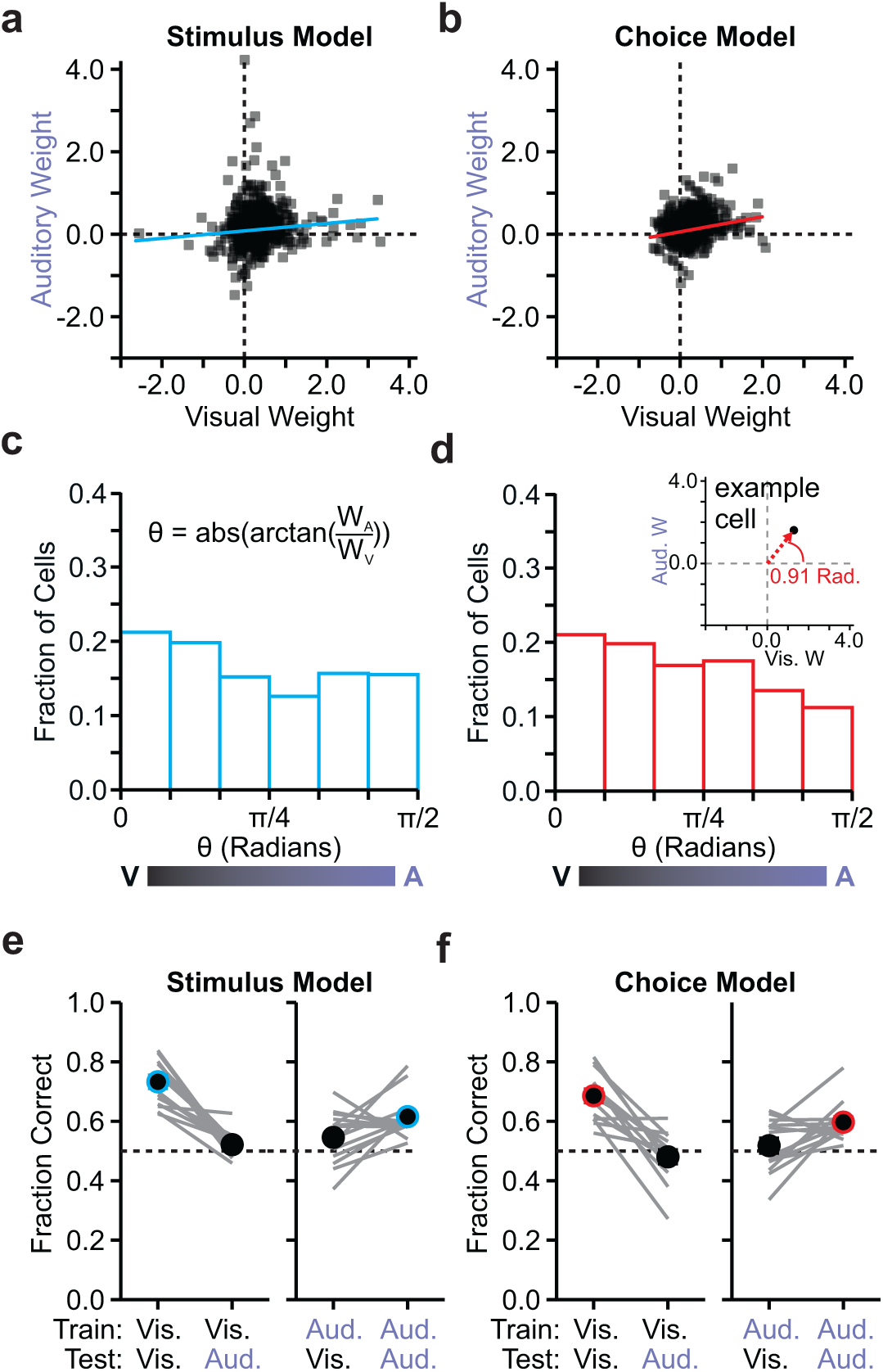
The neural code for stimulus and choice is separable across modalities. **a)** Comparison of weight for each task-responsive neuron (late distractor or target responsive) in the stimulus model when trained with visual or auditory trials. Line is linear fit across all points (R=0.097, p<0.05). **b)** Same as **a** for the choice model (R=0.22,p<0.0001). **c)** Histogram of the angle (θ) between visual and auditory stimulus weights (W_V_ and W_A_), transformed to lie between 0 and 1.6 (π/2) radians (Rad.). Neurons with θ near 0 have larger W_V_, near 1.6 have larger W_A_, and near 0.8 (π/4) have equivalent weights. Inset: equation to calculate θ. **d)** Same as **c** for choice weights. Inset: example neuron weights transformed to θ. **e)** Left: Performance of stimulus model trained with visual trials when tested with visual or auditory trials. Right: Performance of stimulus model trained on auditory trials when tested with visual or auditory trials. Colored circles indicate performance above chance (p<0.0001) and error is S.E.M. across experiments. **f)** Same as **e**, for the choice model. Colored points: p<0.0001.

If the codes are truly separable, then the weights derived from training on one modality should be unable to predict the opposite modality. Indeed, neither model could predict the opposite modality’s stimuli or choices above chance (train visual, test auditory: stimulus – p=0.069, choice – p=0.40; train auditory, test visual: stimulus – p=0.074, choice – p=0.48; Student’s t-test; Figure 5e-f). We also tested the independence of these models by comparing the visual or auditory trained model to a model trained on a combination of auditory and visual trials. The neuronal weights in the single modality models were greater than in the combination models (one-way ANOVA with Tukey’s post-hoc tests; stimulus (p<0.0001): combination vs. all others – p<0.01; choice (p<0.0001): combination vs. visual only – p<0.0001, trend for combination vs. auditory only – p=0.10; Supplemental Figure 5c-f), suggesting that neurons are less predictive in the combination model due to a decrease in signal-to-noise. Taken together, these results demonstrate that visual and auditory stimuli and choices are represented by unique combinations of the populations of V1 neurons.

### Model performance on invalidly-cued trials suggests attention alters decoding and not encoding of visual signals

We find that V1 has separable codes representing both stimuli and choices on visual and auditory trials. Thus, our data are potentially consistent with both proposed hypotheses for how attention alters behavior (i.e. through effects on either representation or read-out). To test whether there are effects of attention on both representation and read-out, we investigated how each of these models performed when tested with the same modality target across attentional states. Specifically, we used invalidly-cued visual trials to test if the models performed differently on expected versus unexpected targets (Figure 6a). There were too few invalidly-cued visual trials to train a new model; instead, we trained the models on stimuli from the validly-cued visual or auditory trials and then tested with stimuli from the invalidly-cued trials.

**Figure 6.**
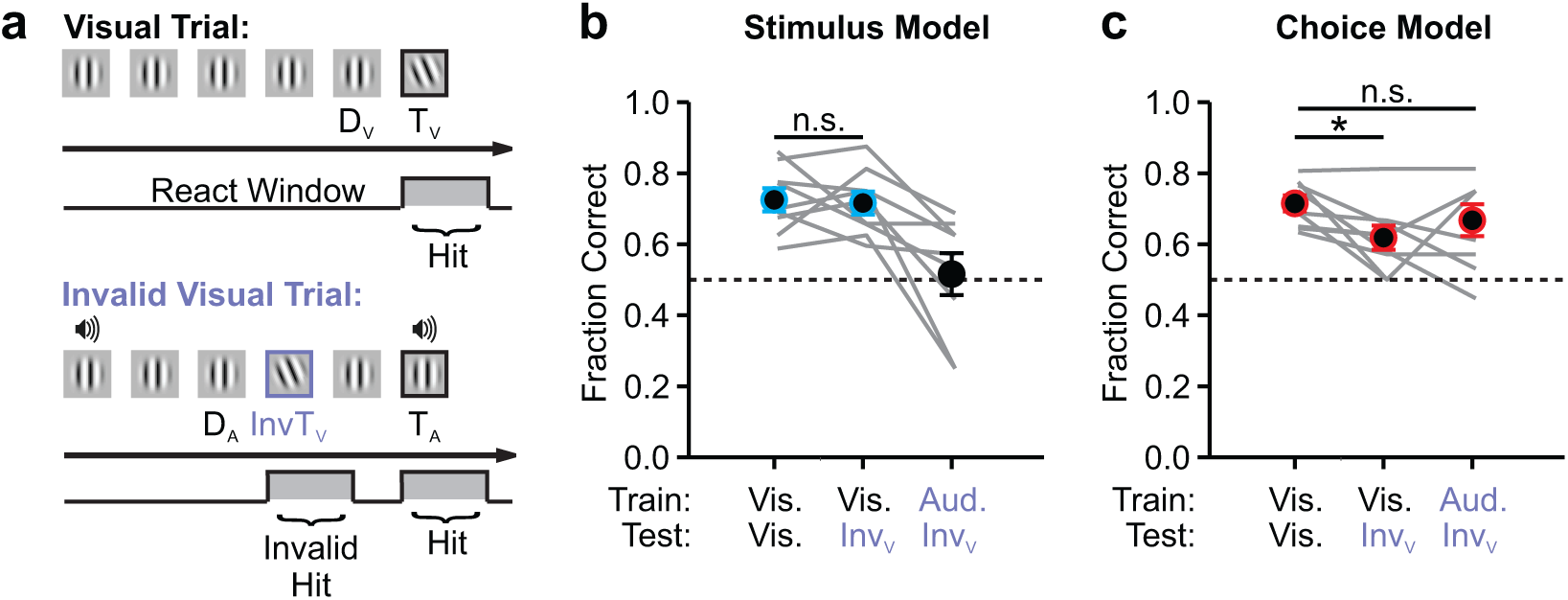
Attention improves the read-out, but not the representation, of visual stimuli in V1. **a)** Schematic of example visual (top) and invalid visual (bottom) trials. **b)** Performance of the stimulus model when trained with either visual or auditory trials and tested with valid (left) or invalid (middle) visual or valid auditory (right) trials. Colored points: p<0.01 compared to chance, Student’s t-test. **c)** Same as **b**, for the choice model; all points: p<0.05 compared to chance. *p<0.05, paired t-test.

When the stimulus model was trained on validly-cued visual trials, the model performed equally well on held-out validly- and invalidly-cued visual stimuli (p=0.82, n=5 mice, 9 sessions; paired t-test; Figure 6b). In comparison, invalidly-cued visual stimuli could not be predicted from a model trained on validly-cued auditory trials (p=0.78, Student’s t-test, Figure 6b). Thus, the stimulus models can only discriminate within-modality stimuli irrespective of attentional state. Moreover, the lack of change in performance on validly-compared to invalidly-cued visual trials suggests that the representation of visual targets and distractors in V1 is not improved with attention.

Unlike the stimulus model, when the choice model was trained on validly-cued visual trials, it was significantly worse at predicting choices on invalidly-cued trials than held-out validly-cued trials (p<0.05, paired t-test; Figure 6c). Yet, when the choice model was trained on validly-cued auditory trials and tested with invalidly-cued visual trials, it performed similarly to the visual model at predicting choices on validly-cued visual trials (p=0.35, paired t-test). Thus, the population code that best discriminates the animal’s choice depends on attentional state, not modality. Moreover, since the representation of choice in V1 depends on the interaction between V1 and its targets, our data suggests that the major effect of attention is to change which population of V1 neurons effectively drives downstream areas to make a decision.

## Discussion

Goal-directed attention changes the activity of sensory neurons, but it is unclear whether these changes in activity relate more to improvements in the encoding of stimuli or to changes in the behavioral read-out of those stimuli. Here, we have developed a task to probe the neuronal correlates of attention in mouse V1. We find that, in addition to altering the magnitude of visually-driven responses of V1 neurons, attention alters how populations in the visual cortex encode choice from trial-to-trial. This supports a model whereby attention acts by changing the population of V1 neurons being read-out by downstream areas.

Our cross-modal detection task was designed to resemble the classic selective attention paradigms typically used in human (Ciaramitaro et al., 2007; Posner et al., 1980) and non-human primate studies (Cohen and Maunsell, 2009; McAdams and Maunsell, 1999; Mehta et al., 2000a). The use of head-fixed mice enables tight control of stimulus presentation and the ability to monitor eye position. In addition, the similarity of task structure across attention conditions rules out the contribution of non-specific cognitive contributions such as changes in arousal, engagement or reward expectation.

Consistent with previous studies (Ciaramitaro et al., 2007; Hembrook-Short et al., 2017; Karns and Knight, 2008; McAdams and Maunsell, 1999; Mehta et al., 2000a), we find that attention towards the visual stimulus increases responses of V1 neurons. There was diversity in the direction of modulation, and neurons that were tuned for task stimuli tended to be more strongly driven on visual trials. Since we could not explore the full tuning curve of neurons across attention conditions, we cannot directly test whether this observed increase in visual responses on visual trials is due to a gain change, as has been seen in other visual attention paradigms (Lee and Maunsell, 2010; McAdams and Maunsell, 1999). However, while task-tuned neurons were more likely to increase their activity with attention, many neurons actually decreased their activity with attention. Further, differences in attentional modulation as a function of orientation preference were due to differences in the number of modulated neurons, not the magnitude of modulation as would be expected from a gain change. Instead, our results support a model whereby attention selectively modulates the activity of informative cells (Hembrook-Short et al., 2017; Verghese et al., 2012). Additionally, tuning-specific modulation of V1 neurons is unlikely to be explained by modulation of the relatively untuned lateral geniculate nucleus of the thalamus, as has been seen in a cross-modal sensory selection task (Nakajima et al., 2019; Wimmer et al., 2015). How the activity of V1 and other areas is modulated by attention may be specific to task-design: unlike the sensory selection task, the mice in our task are incentivized to respond to any target regardless of modality or cue.

The difference in decoding of V1 population activity across task modalities provides additional evidence that neurons are specifically modulated across attention conditions. Consistent with other studies of the relationship between neuronal activity and behavior (Choe et al., 2014; Nienborg et al., 2012; Ruff and Cohen, 2018; Yang et al., 2015), we find that the activity of V1 neurons is tightly linked to both the sensory stimulus and the mouse’s choice. However, while the weights for stimulus and choice within a modality are highly correlated, the weights across modality are nearly uncorrelated. The lack of correlation across stimulus weights is consistent as the sensory stimuli differ across conditions; however, the lack of correlation across choice weights suggests that read-out of V1 activity is changing with attention from trial-to-trial. Indeed, while V1 neurons are equally good at encoding visual stimuli across attention states, they are less good at representing visual choices in the unattended condition. Similar effects of attention on the read-out of visual cortical activity have been suggested to occur in humans and non-human primates (Gregoriou et al., 2009; Pestilli et al., 2011; Ruff and Cohen, 2018), potentially mediated by changes in functional connectivity within (Cohen and Newsome, 2008; Hembrook-Short et al., 2019; Ruff and Cohen, 2014) and between cortical areas (Lakatos et al., 2009; Ruff and Cohen, 2016).

We found a strong representation of auditory stimuli and choices in V1. There is a robust projection from primary auditory cortex to V1 (Charbonneau et al., 2012), and auditory stimuli can modulate visual responses in V1 (Ibrahim et al., 2016; Iurilli et al., 2012; McClure and Polack, 2019). However, our finding that auditory stimuli could only be predicted from V1 in behaving animals has also been reported in humans (Cate et al., 2009; Matusz et al., 2016), making it unlikely that the presence of auditory signals in V1 could be explained by passive sensory transmission. Instead, representation of auditory stimuli in V1 may reflect the presence of auditory choice signals since these are closely related and highly correlated. This is also consistent with the observation that choice-related activity is broadly distributed across the brain (Katz et al., 2016; Pitkow et al., 2015; Runyan et al., 2017). Even so, it is surprising that despite the pervasive representation of choice, visual and auditory choices do not share the same neuronal code. We argue that this separable representation of choice across attentional states reflects differences in the read-out of visual cortical activity. Our data comparing validly and invalidly-cued choices further argue that attention biases downstream areas toward monitoring the most informative V1 neurons.

The reliable changes read-out with attentional state that suggest that modest changes in individual neuron activity can have larger effects on behavior when the entire population is considered. There are a number of functional mechanisms that could explain the observed changes in read-out across attentional states. One possibility is that the attention-dependent changes in V1 activity might alter the feed-forward functional connectivity between V1 and the downstream decoder, thereby engaging separable populations of neurons in the decision (Ruff and Cohen, 2018). Alternatively, cross-modal attention may involve switching between two decoders that each monitor unique populations of V1 neurons. Finally, the observed choice-related activity in V1 may be due to changes in feedback into V1 (Bondy et al., 2018; Yang et al., 2015). Importantly, all of these mechanisms support a model whereby attention alters the activity of specific populations of V1 neurons to enhance the read-out rather than representation of V1 activity.

## Methods

### Animals

All experimental procedures were carried out under a protocol approved by Duke University’s Institutional Animal Care and Use Committee. 17 adult male and female mice were used in this study (>P45, under a regular 12-hour light/dark cycle). All mice used for behavior (n=8) were the F1 offspring of C57/B6J (Jackson Labs #000664) and CBA/CaJ (Jackson Labs #000654). Mice used in naïve experiments (n=9) were either Ai93 (tm93.1(tetO-GCaMP6f)Hze, Jackson Labs #024103; n=5) crossed to EMX1-IRES-Cre (Jackson Labs #005628) and CaMK2a-tTA (Jackson Labs #003010) backcrossed to CBA/CaJ (25-45% CBA), or the F1 offspring of C57/B6J and CBA/CaJ (n=4). Ai93 mice were fed Doxycycline chow (200 mg/mL) (from onset of pregnancy until postnatal day 45 (P45)) to suppress calcium indicator expression and decrease the likelihood of seizures (Steinmetz et al., 2017).

### Cranial window implant

Mice were implanted with chronic cranial windows as previously described (Goldey et al., 2014). Prior to surgery (3-16 hours), mice were injected with dexamethasone (3.2 mg/kg, subcutaneously (SC), Bimedia) to reduce brain swelling during the craniotomy. Immediately before surgery, mice were given prophylactic analgesia (2.5 mg/kg meloxicam, SC) and anesthesia was induced with a combination of ketamine and xylazine (200 mg/kg and 30 mg/kg, intraperitoneally (IP)) and 4% isoflurane. Stable anesthesia was maintained at 1-1.5% isoflurane for the duration of the surgery. A titanium headplate was attached to the skull with dental cement (C&B Metabond, Parkell) and a 5 mm craniotomy was drilled centered on the left visual cortex (3.1 mm lateral and 1.6 mm anterior from lambda). The craniotomy was sealed with a glass window (an 8 mm coverslip bonded to two 5 mm coverslips (Warner no. 1) using a refractive index matched adhesive (no. 17, Norland)) using dental cement. After surgery, mice were recovered on a warm heating pad and given analgesics (buprenorphine, 0.5 mg/kg, SC) and antibiotics (cefazolin, 50 mg/kg, SC) for 48 hours.

### Sensory Stimulation

Visual stimuli were presented on a calibrated (i1 Display Pro, X-Rite) 144 Hz LCD monitor (Asus) placed 21 cm from the right eye (contralateral to the craniotomy) perpendicular to the mouse. All visual stimuli during the behavioral task were static, sinusoidal gabor patches (30-50° diameter, 0.1 cycles per degree, 100% contrast, 60 cd/m^2^ mean luminance). Auditory stimuli were either pure tones (task cue and target stimuli), white noise (feedback on error trials), or multiple tones (feedback on correct trials) and were delivered via speakers placed behind the mouse (max amplitude ∼90 decibels). After each behavioral session, drifting gratings (2 Hz, 8 or 16 directions in 45° or 22.5° increments) were presented to the passively viewing mouse at the same position, size and spatial frequency as the task stimuli. All sensory stimuli were delivered, and synced to imaging acquisition when applicable, via custom software created in MWorks (http://mworks-project.org).

### Retinotopic Mapping

After at least 1 week of recovery from surgery and habituation to head restraint, visual cortex was retinotopically mapped by wide-field imaging of intrinsic autofluorescence or GCaMP signals through the cranial window (Andermann et al., 2011). While head-restrained on a running wheel, mice passively viewed vertical (0°) drifting gratings at 2-4 retinotopic locations (30° diameter; 5° and 35° in azimuth and either 15° or ±15° elevation). For intrinsic autofluorescence imaging, stimuli were presented for 10 s, with 10 s of mean luminance between each presentation. Changes fluorescence were monitored by illuminating the cortex with blue light (white light (Exfo) or 473 nm LED (Thorlabs) with a 462±15 nm band filter (Edmund Optics)) and collecting emitted green and red light (500 nm longpass filter), monitored with a CCD camera (Rolera EMC-2, Qimaging) at 2 Hz through a 5x air immersion objective (0.14 numerical aperture (NA), Mitutoyo), using Micromanager acquisition software (NIH). Visually-driven changes in cortical activity were measured by calculating the normalized change in fluorescence (dF/F), where F is the average fluorescence of the whole movie, during stimulus presentation for each position. For GCaMP imaging, the setup was the same, except the stimulus was presented for 5 s, with a 5 s inter-stimulus interval, and the emitted light was collected via a green bandpass filter (530±15 nm, Edmund Optics).

### Virus Expression

For all behaving mice, we targeted injections of AAV1.Syn.GCaMP6m.WPRE.SV40 (titer: 1.1-2.2×10^13^ GC/ml) into lateral V1 using the intrinsic signal retinotopic map and vasculature pattern as a guide. Naïve wildtype mice received injections of GCaMP6m, or AAV1.Syn.NES-jRGECO1a.WPRE.AV40 (titer: ∼6.5×10^12^ GC/ml) into V1. Virus was diluted 3:1 with Texas Red dye (10 mM in saline, Life Technologies) and loaded into a glass pipette (World Precision Instruments (WPI)) with a broken, beveled tip ∼20 µm in diameter. The pipette was inserted into a Hamilton syringe which was mounted in a syringe pump (WPI). Following removal of the glass window, the pipette was lowered into the craniotomy and 100 nl of virus was injected at two depths (250 and 450 μm) at a rate of 100 nl/minute. The pipette was left in the tissue for 5-10 minutes and the dye was visualized to check for diffusion into the tissue. Finally, a new glass window was replaced into the craniotomy and sealed with cement.

### Behavioral task

All behaving mice were either water (n=7 mice) or food (n=1) scheduled. Water scheduled mice received 0.1M saccharine water (Acros Organics) and food scheduled mice received liquid nutritional shake as reward (Ensure, vanilla flavor). Mice were supplemented with plain water or food pellets if they did not receive all of their allotted water or calories for the day during training. The behavior training and testing occurred during the light cycle.

Mice were trained to perform the cross-modal detection task in the following steps. On the first day of training, mice were head-restrained in a custom-built behavioral rig (parts from Thorlabs, Newport, Digikey and Standa (Histed et al., 2012)). To earn reward, mice were required to press and hold the lever for at least 400 ms and no longer than 10 s. At the time of lever press, a 6 kHz tone was played (this would later be the cue for auditory trials), and if the mouse continued to hold the lever for 400 ms, a 10 kHz target tone was played indicating the onset of the reaction window. After the target presentation, mice were allowed up to 10 seconds to release the lever. Releasing the lever within this window resulted in reward delivery and auditory feedback indicating a correct response. Releasing the lever before the target tone (early release), or failure to release the lever within the reaction window (miss) resulted in auditory feedback (white noise) indicating an error and a timeout (1-6 s). Each trial was interleaved with inter-trial interval (ITI; 4-6 s) during which a new trial could not be initiated. Once mice began to reliably release the lever after the target tone we followed several steps to gradually make the task more difficult, roughly in chronological order: 1) increasing the random delay between the cue and the target up to four seconds so that the mice could not use a timing strategy to detect target tones 2) decreasing the allowed reaction time from 10 seconds to 550 ms, and 3) adding more difficult targets (lower amplitudes) around the animal’s threshold. Mice were considered to have learned the auditory task if they performed better than 90% correct on easy targets. This paradigm was continued for two to three weeks then the mouse was switched to learning the visual task in a similarly structured paradigm (even if the mouse was not yet fully trained on the auditory task).

While mice already knew how to use the lever to earn reward, all mice needed to be retrained to detect visual stimuli, suggesting that they do not generalize across modalities. Thus, we used the same paradigm to train the mice to detect target gratings as we had for target tones. On the first day of visual training, a full-field, vertical grating appeared upon trial initiation. If the mouse held the lever for 400 ms, the grating changed 90° to a target orientation. The target grating then stayed on the screen until the mouse either released the lever or the reaction window ended. Mice typically began reliably releasing the lever during the target stimulus within approximately five sessions. To make the task more difficult we gradually 1) increased the random required hold time, 2) decreased the reaction time, 3) decreased the size of the stimulus, 4) moved the stimulus to the right (to be closer to the retinotopic location of the future injection site), 5) added a mean luminance inter-stimulus interval (ISI) during the anticipation period to mask the motion signal in the transition from distractor to target, and finally, 6) increased the difficulty by reducing the difference between the distractor (0°, vertical grating) and targets (any stimulus that is not vertical).

After the mice were proficient at the visual task, they were trained on the visual and auditory tasks on interleaved days until they consistently 1) got at least 90% correct on the easiest trials, and 2) less than 50% of trials were early releases. Finally, we randomly interleaved visual and auditory trials within the same session. At this point, the visual distractor stimulus was added to the auditory trials.

In the final form of the task (Figure 1a-c), each trial was initiated when the ITI ended and the mouse had pressed the lever. The trial start triggered the presentation of a 100 ms, vertical, sinusoidal gabor patch (30° or 50° in diameter, 15 to 30° in azimuth, 0° in elevation; one mouse had a 200 ms stimulus for some sessions) followed by a 250 ms ISI. On each trial, a target was presented after a variable number of distractor presentations (2-10, flat distribution). On auditory trials, the first visual distractor stimulus was paired with a 6 kHz tone which cued the mouse to expect an auditory target (a 10 kHz tone paired with a visual distractor stimulus). The absence of a tone on the first distractor cued the mouse to expect a visual target (any non-vertical stimulus). Mice received reward if they released the lever within 100-550 ms (sometimes extended to 1000 ms) after a target occurred. For behavioral and neuronal analyses, a narrower reaction window (visual: 200-550 ms, auditory: 150-450 ms) was used to ensure that the majority of releases in this window were due to stimulus-driven behaviors and have independent reaction windows for each stimulus presentation within the trial.

Invalidly cued visual or auditory targets (Figure 1d-g, Figure 6), in which the trial was cued as one modality but the target delivered was of the opposite modality, were delivered on 2.4± 0.13% of trials. Invalidly cued targets could appear after 1-9 distractor presentations, as they always appeared between the cue and the validly cued target. In the case that the mouse failed to respond to an invalidly cued target, the trial continued and the mouse had the opportunity to detect a validly cued target. For analyses where attention was tested across valid and invalidly cued trials (Figure 1d-f), all invalid hits were rewarded.

Two of the 5 mice with attention were trained and imaged without rewarding invalid hits and were tested with rewarded invalid trials in later sessions. These mice had a greater effect of attention when tested with rewarded invalid trials (hit rate across mice on invalid trials normalized to valid trials: training rewarded – 79.5±0.2%, training not rewarded – 57.2±5%, mean ± S.E.M across experiments). However, the fraction of attention modulated V1 neurons was similar for these two groups (fraction of late responsive neurons significantly modulated by attention: training rewarded – 0.16±0.04, training not rewarded – 0.15±0.04, mean ± S.E.M across experiments). Thus, we considered all 5 mice together for imaging analyses.

During imaging sessions, the position of the visual stimulus was optimized to best activate the imaged neurons by performing a brief retinotopy experiment at the beginning of each session. The same position was used for both the behavior and passive tuning experiments. Behavior sessions during imaging were on average 49±3 minutes (range: 30-60 minutes, 2-4 sessions per mouse, 309±20 (range: 180-434) trials per session). Naïve imaging sessions were on average 53±3 minutes (range: 30-66 minutes, 1-3 sessions per mouse, 379±30 trials (range: 190-500)).

### Two-photon Imaging

Fluorescence of genetically encoded calcium indicators was monitored in populations of neurons with a two-photon microscope (Neurolabware) and collected with Scanbox acquisition software. Excitation laser light (Mai Tai eHP DeepSee, Newport; tuned to 920 nm for GCaMP or 1020-1040 nm for jRGECO) was raster scanned with a resonant galvonometer (8 kHz, Cambridge Technologies) onto the brain via a 16x or 25x (0.8 or 1.1 NA, Nikon) water-immersion objective into a rectangular plane 582±54 by 273±34 μm in size (X by Y; range: X: 278-1030 μm, Y: 117-581 μm) and 257±7 μm below the pial surface (range: 198-303 μm). Laser power out of the objective ranged from 30 to 80 mW. Emitted light was passed to a dichroic mirror (562 nm cut-off (Semrock)) and directed toward GaAsP photomultiplier tubes (H10770B-40, Hamamatsu) via either a green (510±42 nm (Semrock)) or red (607±35 nm (Semrock)) bandpass filter. Images were acquired at 30 Hz and aligned to behavioral and visual variables.

### Pupil Imaging

Partially scattered infrared light from the two-photon excitation was emitted from the pupil and collected with a Genie Nano CMOS (Teledyne DALSA) camera using a longpass filter (695 nm) at 30 Hz. Pupil data was collected simultaneously with two-photon imaging in three mice; for two mice, pupil data was collected in a separate set of experiments.

### Data Processing and Analysis

All analyses were performed with custom code written in MATLAB (Mathworks).

#### Behavior

Behavioral sessions were manually cropped to include only stable periods of performance by removing periods within a session with high lapse rates (misses on the easiest target conditions) or early release rates (lever releases before the target appears). Sessions included for analysis were further restricted to have 1) at least 90% correct on one of the two easiest levels on both auditory and visual trials, 2) at most 35% of trials be early releases. Thus, 29±4 sessions (range: 8-40) and 7586±1610 trials (range: 2043-13203) were included for each mouse.

Each stimulus presentation following the 2^nd^ distractor in each trial was categorized as either a hit, miss, false alarm (FA), or correct reject (CR) based on the time of release relative to a target or distractor onset. Lever releases between 200 and 550 ms after a target on visual trials and 150 and 450 ms after a target on auditory trials were hits. Conversely, failure to release by the end of these windows was considered a miss. Releases, or failure to release, during similar windows following a distractor was considered a FA or a CR.

Behavioral performance was primarily analyzed by pooling across all test sessions (Figure 1, Supplemental Figures 6, 2) and but also evaluated on a session by session basis (Supplemental Figure 1) to measure the effect of the cue on performance. To account for small differences in target difficulty levels used for each session, targets were binned into six logarithmically spaced groups that spanned the minimum and maximum target values (visual: 8° to 90°, auditory: 0.03 to 100% of max amplitude). Hit rate was computed from each session from the number of hits and misses for each target type:

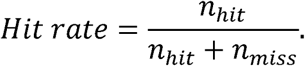

Lapse rate was measured as 1 – Hit rate for the easiest target of a session (within each modality) and FA rate was computed from each session from the number of FAs and CRs (Figure 1g):

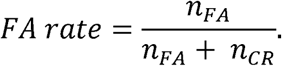

Within each modality, hit rates across target difficulties and the FA rate (representing a 0° or 0% amplitude target) were fit with a Weibull function to determine discrimination thresholds (50% of the upper asymptote to account for lapse rate).

Reaction time was calculated as the mean time of lever release from target onset for the target type in question (Supplemental Figure 6). We did not observe any consistent effect of the cue on reaction time (validly-cued vs invalidly-cued - visual: p=0.39; auditory: p=0.07; paired t-test; Supplemental Figure 6a-b). However, since only visual trial reaction times depended on trial difficultly (visual: p<0.01; auditory: p=0.48; one-way ANOVA, Supplemental Figure 6c-e), we may be limited in our ability to resolve reaction time differences across attentional states.

To test attention toward the cued modality we compared each mouse’s response to validly and invalidly cued targets. Mice were considered to have an effect of attention if the hit rate on validly cued targets was statistically greater than hit rate on invalidly cued targets (matched for target difficulty) across both modalities (Figure 1f**, left**). To match target types, the proportion of each invalid target type (by difficulty and modality type) was determined and valid trials were randomly subsampled for each stimulus type to match that distribution.

#### Pupil tracking

The size and position of the pupil was extracted from each frame using the native MATLAB function *imfindcircles*. Pupil size was quantified as the fraction of change from baseline (one second before trial start) then re-normalized by subtraction relative to the start of each trial or target presentation. Pupil position was quantified as the change in the horizontal and vertical position of the center of the pupil from baseline (one second before the start of each trial or target presentation). Analysis windows were chosen to match two-photon imaging analysis windows during the anticipation phase of the trial (1400-2833 ms after the start) or after the target (100-200 ms after the target). Changes in pupil position were converted to degrees of visual angle with a 1:25 degrees to μm scale (Park et al., 2012).

#### Two-photon Imaging

### Registration and segmentation

Image stacks were registered for x-y motion to one stable, 100-frame average reference image, using Fourier domain subpixel 2D rigid body registration. dF/F was calculated on a trial-by-trial basis by defining F as the average fluorescence one second prior to trial start. Maximum (Max) dF/F images were found by finding the maximum pixel value across specific task windows (trial start, late anticipation, or target aligned) or passive direction stimulus types. Max dF/F images were used to manually segment and create masks of cell body ROIs. The same masks were used for the behavior and passive tuning experiments.

Pixels within a cell mask were averaged for each registered frame to get the time-course of activity for each cell. Neuropil contamination for each ROI was calculated by first creating a buffer ring of 4 pixels around each cell body, creating a neuropil ring 6 pixels around the buffer that excluded other ROIs, estimating the scaling factor (by maximizing the skew of the subtraction between the cell and neuropil time-course), and finally, subtracting the weighted neuropil time-course from the cell’s time-course. Finally, remaining contamination from brain motion was removed by discarding trials with large, fast changes in dF/F across all cells, which could only be due to changes in the imaging plane and not task-driven neuronal responses.

### Passive orientation tuning

We generated orientation tuning curves for each cell from responses to passively viewed drifting grating. Single trial responses for each cell were measured as the mean dF/F 0-1333 ms after stimulus onset. Stimuli moving in opposite directions were treated as the same orientation, and average responses to each orientation were found for each cell. Responses below zero were set to zero and these response distributions were fit with a von Mises function to get orientation tuning curves

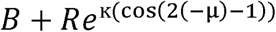

where *B* is the baseline response, *R* is the modulation rate, κ is the concentration, and μ is the preferred orientation.

To determine the reliability of this tuning we bootstrapped the fit by resampling trials 1000 times. A cell was considered to be reliably tuned if the resampled peak of the fit was within 30° of the actual fit 90% of the time. Tuned cells were then grouped into four orientation preference bins (0°, 45°, 90°, and 135°) by finding the closest orientation to the cell’s fit peak.

### Task neuronal activity

Short-latency, visually-evoked responses to task stimuli (i.e. first distractor, late distractor (5th-10th), or target) were measured as the average response 100-200 ms after stimulus onset. Long-latency, visually-evoked responses to the anticipation period were measured as the average response 1400-2833 ms after trial start on trials with at least 8 distractor stimuli. Cells were considered significantly responsive if the mean dF/F during the response window was statistically greater than baseline window (-33-67 ms before the task event) using a one-sided paired t-test across trials. Cells were considered target responsive if their response to either easy (>32°), hard (8°-32°), or all targets was significantly greater than baseline (Bonferroni corrected paired t-test). Cells were considered suppressed if the average response 1400-2833 ms during the anticipation phase was significantly below baseline. Both visual and auditory trials were used to find distractor responsive and suppressed cells, whereas only visual trials were used to find target responsive cells. All analyses were performed on responses from hit or miss trials.

Attention modulation was measured as a selectivity index (SI) between visual and auditory trials for late distractor responsive neurons (Poort et al., 2015). The difference in mean response (*R^-^*) between visual (*V*) and auditory (*A*) trials was normalized by the standard deviation of pooled visual and auditory responses (*S_VA_*) to account for variable responses across trials, where:

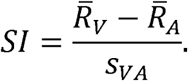

Cells were considered significantly modulated if their SI was consistently positive or negative across 95% of bootstrapped SI (Supplemental Figure 3a). For suppressed cells attention modulation was calculated during a late window of the anticipation phase (1400-2833 ms after trial start).

### Modeling

Each experiment was considered separately for the predicting either the stimulus presented (stimulus model) or animal’s choice to hold or release the lever (choice model) from the single trial responses of simultaneously recorded populations of neurons. For each experiment both models were fit with the same data: the responses of a population of neurons to distractor and target presentations. The only difference between the two model types was how each stimulus presentation was labeled (stimulus model: target or distractor, choice model: yes or no). To process and select the data that went into the models, the following steps were taken. First, all stimulus responses were z-scored. Next, to reduce bias toward representation of one stimulus type in the response distributions we balanced the number of target and distractor stimuli in each model by random selection – responses used were 50% distractors, 25% hard targets, and 25% easy targets (in many datasets, there were no hard auditory trials and therefore we selected 50% targets and distractors). Finally, to avoid over-parameterizing the model, we limited the number of neurons used (maximum 15 neurons). We specifically selected neurons that were either target or late distractor responsive and sharply tuned (90% of the resampled estimates of preferred orientation were within 11.25° of the original estimate). If more than 15 neurons met these criteria, 15 neurons were randomly selected (average number of neurons per dataset: 13±1, range: 8-15). Three naïve experiments did not have enough neurons under these conditions and therefore the tuning criteria was dropped. However, differences in tuning could not explain the differences we observed between behaving and naïve models for predicting hard targets (Figure 4d-e; average performance of experiments that pass strict criteria: 44±2%, p=0.99, n=8 experiments, Student’s t-test). Three other naïve experiments did not have enough task-responsive neurons and therefore were not included in this analysis (minimum 8 neurons). To train the combination-modality model we randomly selected half of the visual trials and half of the auditory trials (Supplemental Figure 5c-f).

Using the stimulus responses of these simultaneously recorded neurons, we trained a logistic regression to discriminate between task stimuli or choices (using MATLAB’s *glmfit* routine) and extracted a weight for each neuron. Fraction correct was determined by applying the neuronal weights from each model to previously untrained population responses from the same neurons. Performance of the within-modality models was tested by performing a hold-one-out analysis across all selected trials used in that model. Cross-modality model tests were performed with all selected trials from the opposite modality. No invalidly-cued stimuli were used to train the models and could thus be directly tested. Finally, the combination-modality model was tested with the half of trials of each modality that was left out of the model fitting procedure. Decision criteria were calculated for each experiment as the fraction of trials of the predicted variable (e.g. the stimulus model decision criterion was 0.5 for each experiment since half of the trials were targets).

To calculate a stimulus and choice weight for each neuron in the dataset, we took a bootstrapping approach. For each experiment, we randomly sampled 15 neurons and calculated their weight for stimulus or choice 1000 times. Thus, each neuron was sampled 153±2 times and the average bootstrapped weight was used for analyses in Figure 5a-b and Supplemental Figure 5e-f.

### Experimental Design and Statistical Analysis

Sample sizes were not predetermined by statistical methods, but our sample sizes of the neurons and animals are similar to other studies. The numbers of cells, animals or experiments are provided in the corresponding text, figures and figure legends. All error values in the text are standard error of the mean and all tests for significance are two-tailed unless otherwise specified. Data collection and analysis were not performed blind to experimental conditions, but all visual presentation conditions were randomized.

### Data and code availability

Data will be made available by reasonable request to the corresponding author. Custom code written in MATLAB for data analysis is available on Github: https://github.com/Glickfeld-And-Hull-Laboratories/Manuscripts/CrossModalAttentionV1

## Acknowledgements

We thank E. Burke, C. Dobrott, M. Fowler, J. Issac, and K. Murgas for surgical assistance and A. Yan for assistance with behavioral training. We thank Drs. Fan Wang, Court Hull and Stephen Lisberger for comments on the manuscript and members of the Hull and Glickfeld labs for helpful discussions. This work was supported by grants from the Pew Biomedical Trusts (L.L.G.), the Alfred P. Sloan Foundation (L.L.G), and the NIH: Director’s New Innovator Award (DP2-EY025439, L.L.G.) and Ruth L. Kirschstein Pre-Doctoral Fellowship (F31-EY028018-2, A.M.W.).

## Author contributions

A.M.W. and L.L.G. designed the experiments. A.M.W., J.M.B and L.L.G designed the analysis. A.M.W. collected and analyzed the data. A.M.W. and L.L.G. wrote the manuscript.

---

**Supplemental Figure 1.**
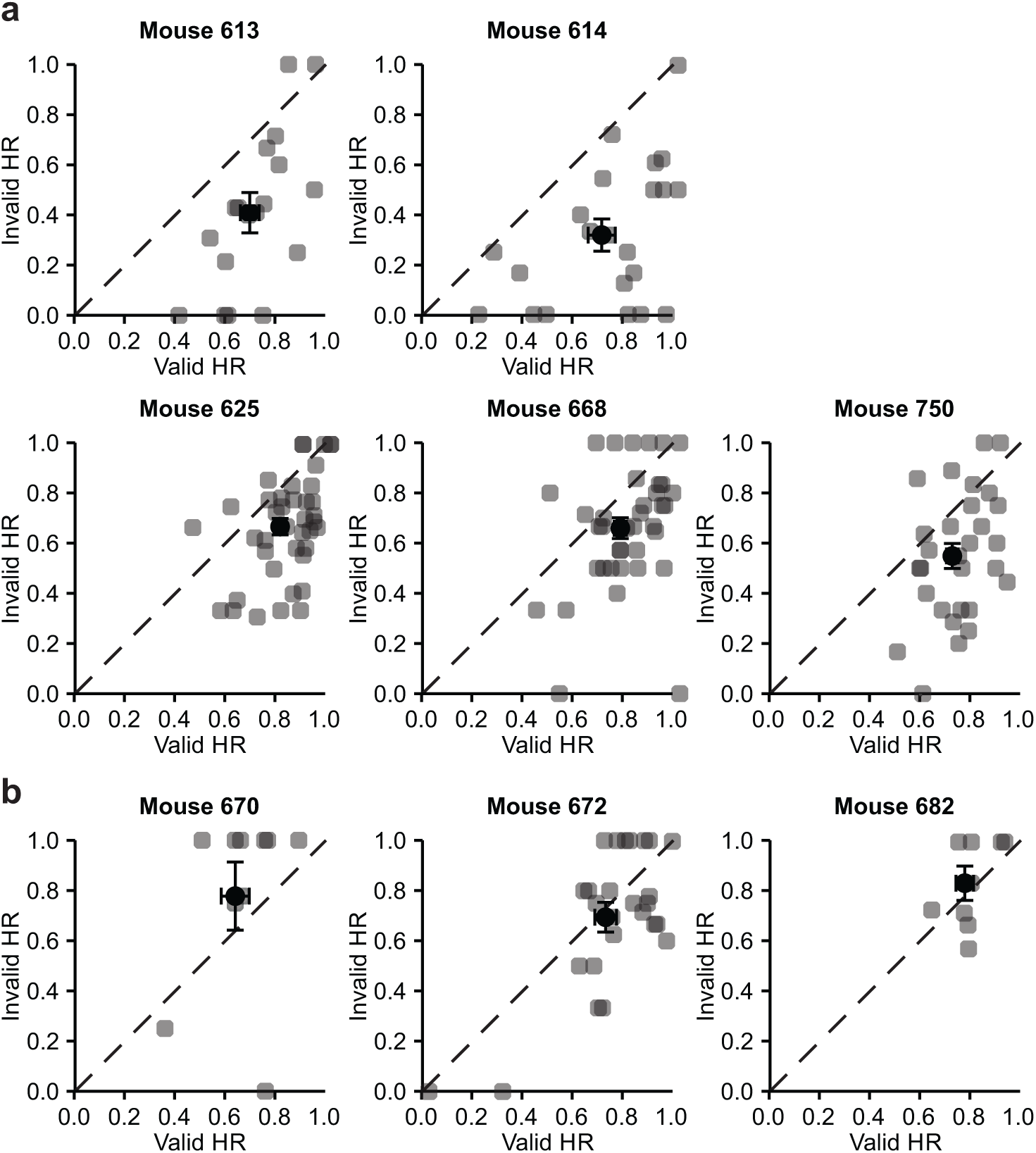
Mice used in imaging analysis have attention effect across sessions. **a)** Hit rate (HR) for each session (gray) and average across sessions (black) for matched valid and invalid trials for each mouse with attention; p<0.001, paired t-test. Mouse 668 is the same mouse shown in Figure 1 d-e. Error is S.E.M. across sessions. **b)** Same as **a**, for mice that did not have a significant effect of attention; 670: p=0.72, 672: p=0.20, 682: p=0.89.

**Supplemental Figure 2.**
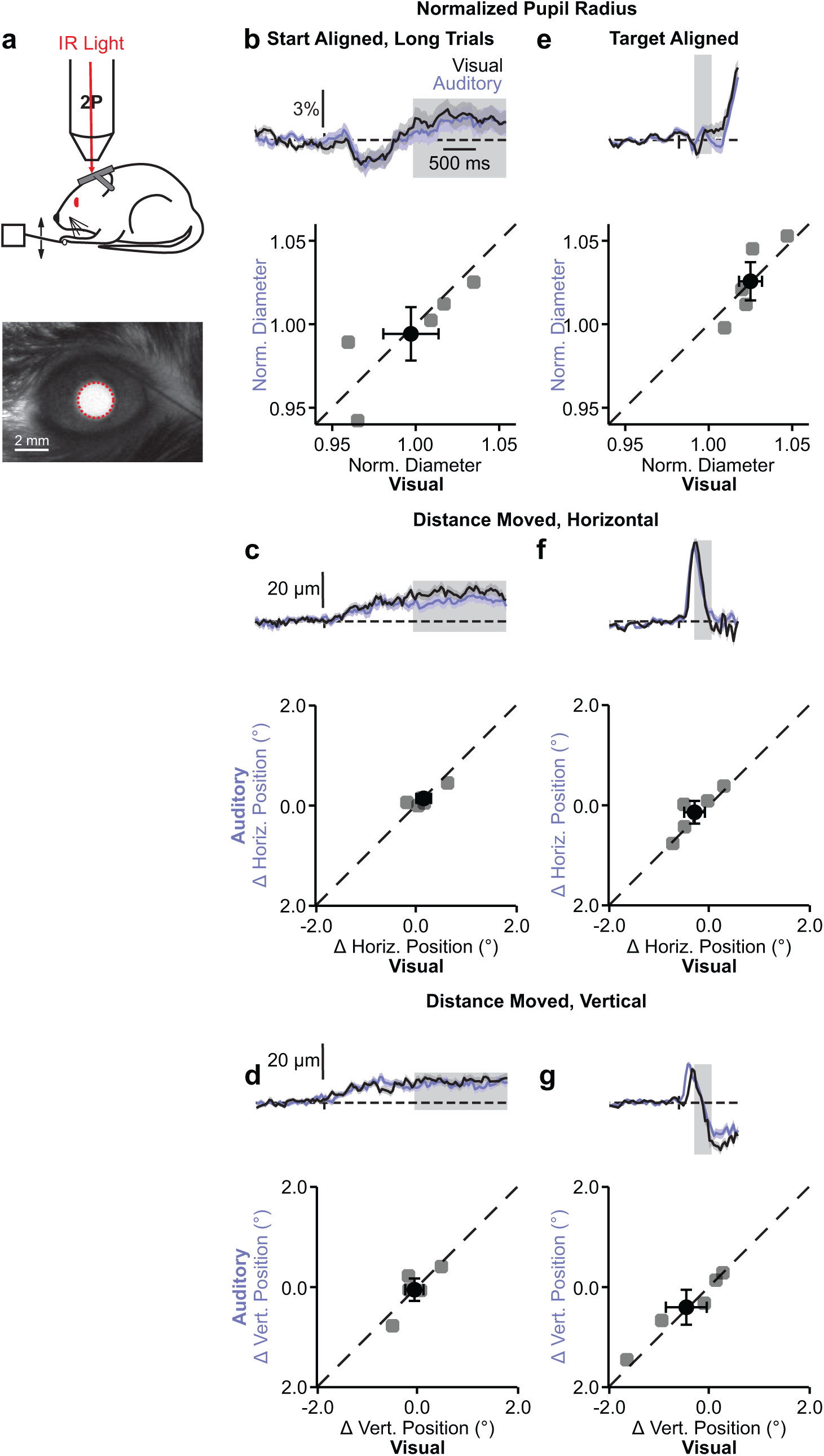
Pupil size and position do not vary across visual and auditory trials. **a)** Top: Schematic of infrared (IR) illumination of the pupil via two-photon excitation; bottom: IR image from example session with tracked pupil outlined (red). **b)** Top: Pupil radius from example session across visual and auditory trials aligned to the trial start. Shaded gray region is the analysis window during the late anticipation period in Figure 2d-e. Bottom: Average normalized pupil diameter during the shaded analysis window for individual mice (gray) and across all mice (black); p=0.80, n = 5 mice, paired t-test. Error is S.E.M. across mice. **c-d)** Same as **b**, for change in horizontal (**c**, p=0.84) or vertical (**d**, p=0.99) pupil position. **e-g)** Same as **b**-**d** aligned to trial target (**e**, p=0.90; **f**, p=0.20; **g**, p=0.65).

**Supplemental Figure 3.**
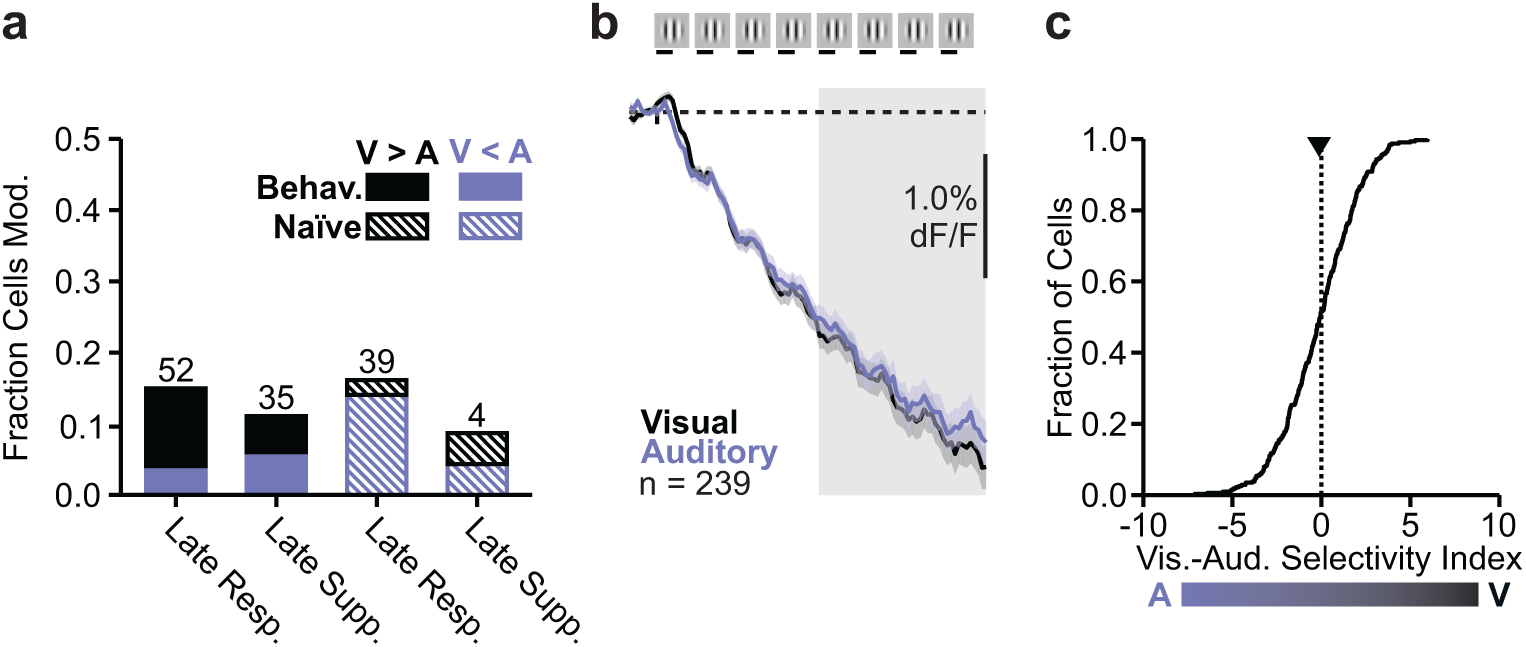
Attentional modulation of responsive or suppressed neurons in behaving and naive mice. **a)** Fraction of neurons that were significantly positively (black) or negatively (purple) modulated on visual trials in behaving (solid) and naïve (dashed) mice. **b)** Average time-course of suppressed neurons on visual (black) and auditory (purple) trials aligned to trial start for trials with at least 8 distractor stimuli. Only neurons from experiments with 100 ms stimuli included. Response late in anticipation period (gray shaded region) is not significantly different across visual and auditory trials (p=0.37, paired t-test). Error is S.E.M. across cells. **c)** Cumulative distribution of SI of suppressed neurons across late trial window. Average SI (filled triangle) is not significantly different from zero (p=0.31, n=316, Student’s t-test).

**Supplemental Figure 4.**
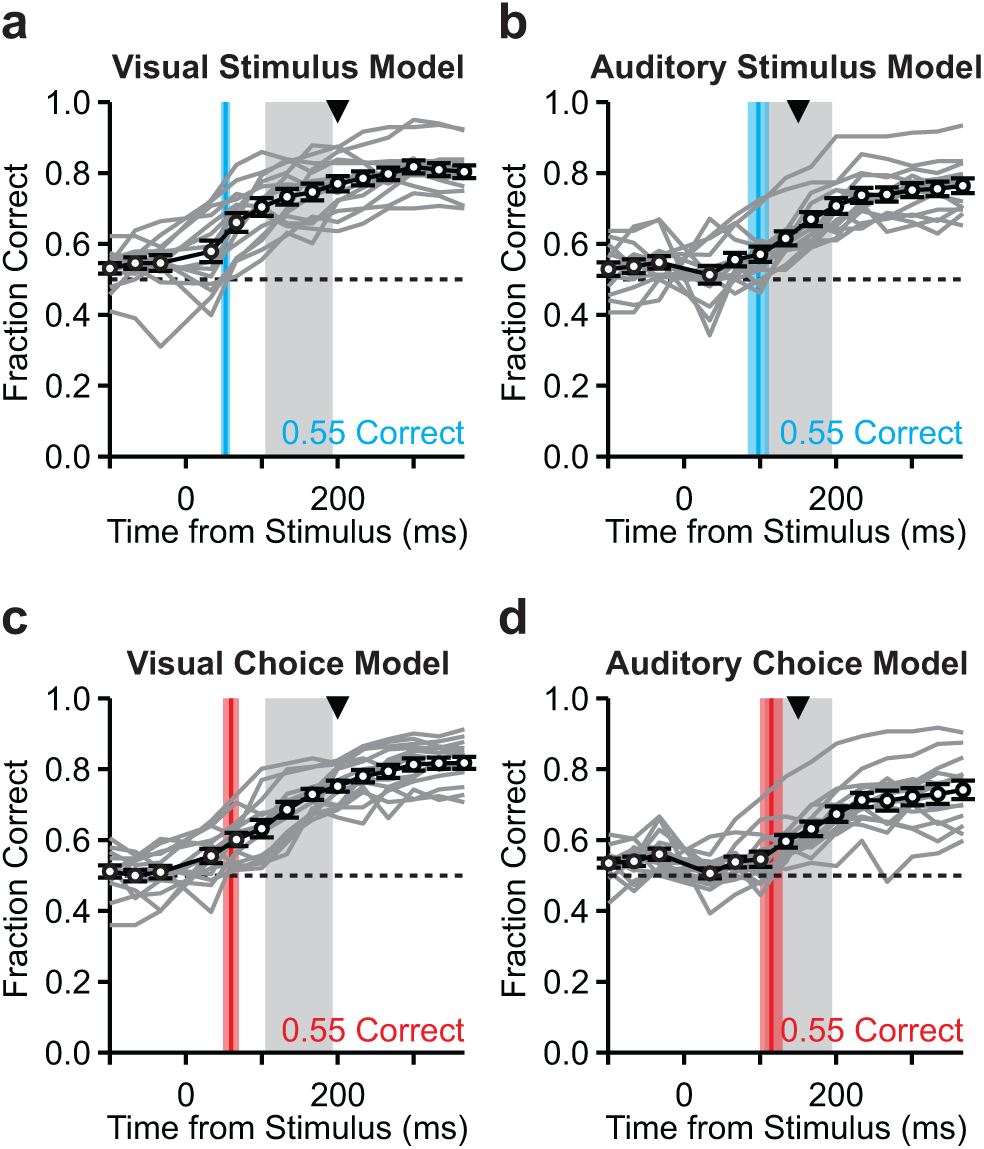
Stimuli and choices can be predicted before rewards occur in both visual and auditory models. **a)** Performance of the stimulus model trained on visual trials for different 100 ms response windows relative to the time the stimulus comes on the screen. Each gray line is one experiment; black is average and error is S.E.M. across experiments. Gray shaded region indicates the window used for all other analyses. Triangle indicates minimum reaction time on visual trials. Blue line and shaded region indicates average time from target when the model performed better than 0.55. **b)** Same as **a**, for the auditory stimulus model. **c-d)** Same as **a**-**b**, for the choice models.

**Supplemental Figure 5.**
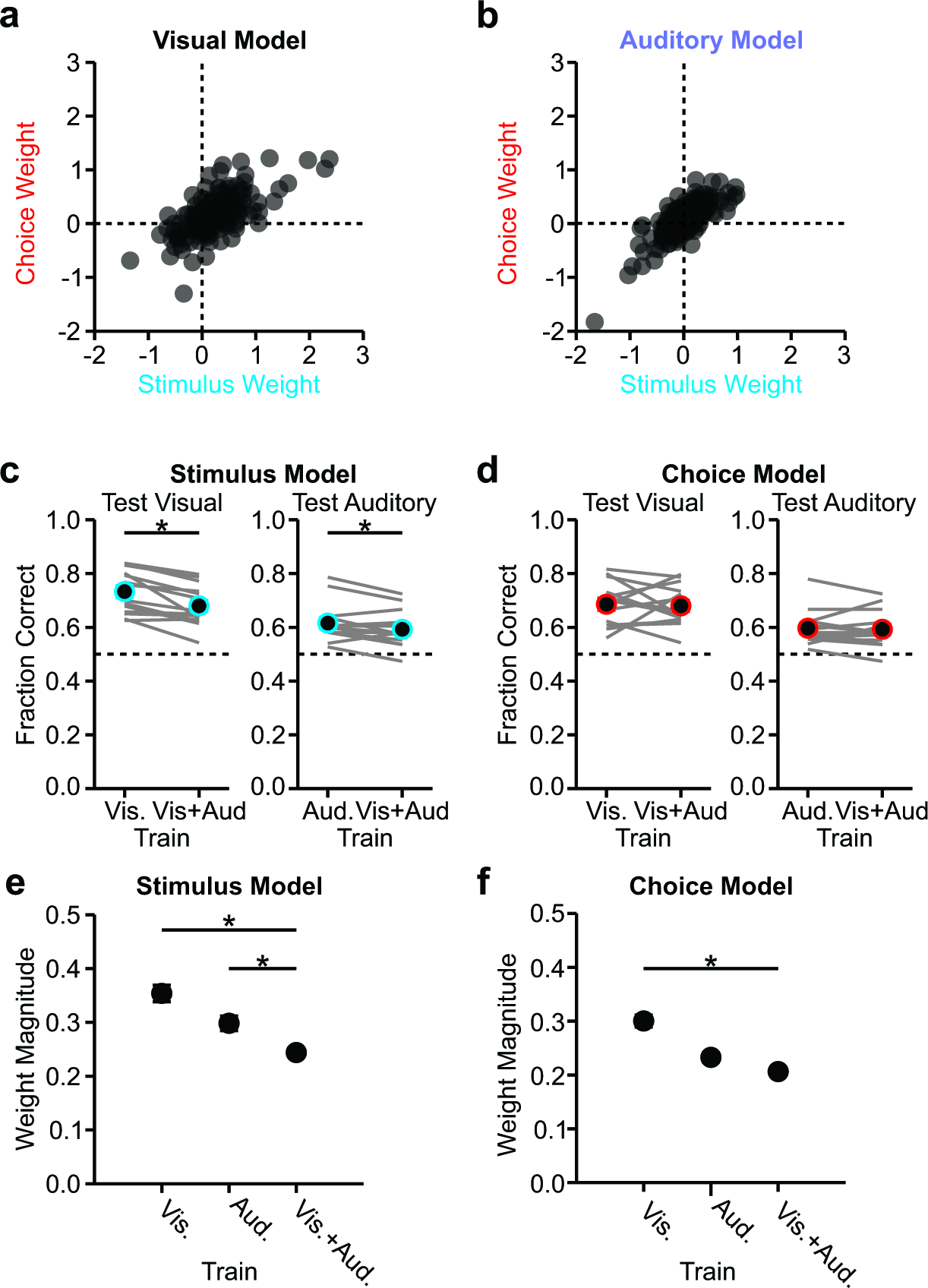
Stimulus and choice weights are correlated within single-modality models, but are smaller in a combination model. **a)** Visual stimulus and choice weights for all late-distractor responsive neurons (determined by bootstrapping model, R=0.63, p<0.0001). **b)** Same as **a**, for auditory weights (R=0.80, p<0.0001). **c)** Stimulus model was trained with a single modality or both modalities together and tested on visual (left) or auditory (right) trials. Each gray line is one experiment. Colored circles are above chance; Error is S.E.M. across sessions; p<0.05, paired t-test. **d)** Same as **c**, for the choice model. **e)** Weight magnitude (absolute value of the bootstrapped weight for each neuron) for visual (left), auditory (middle) and combination (right) stimulus models. One-way ANOVA with Tukey’s post-hoc tests, p<0.01. **f)** Same as **e**, for the choice models. p<0.0001.

**Supplemental Figure 6.**
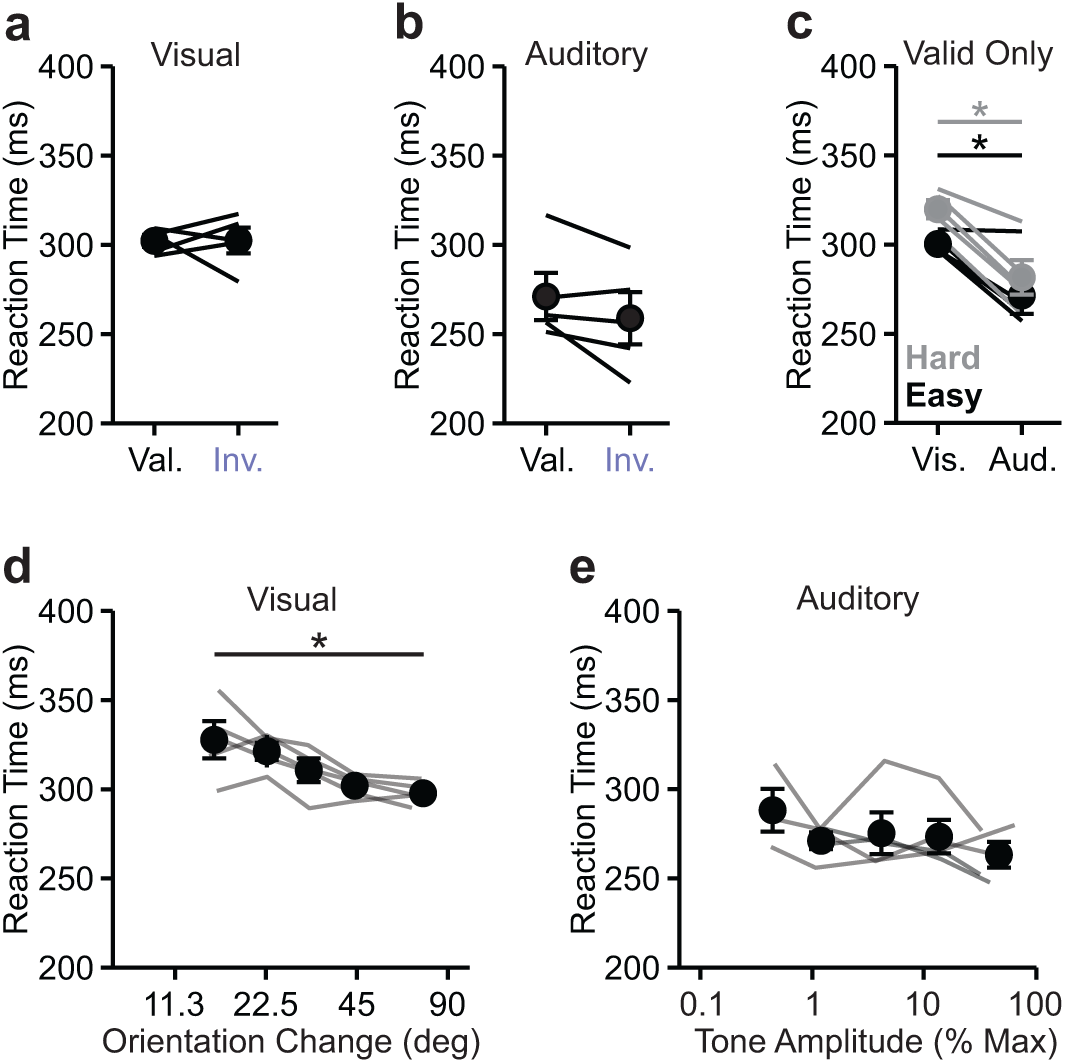
Attention does not affect reaction time. **a)** Reaction times for mice with attention across valid and invalid visual trials matched for difficulty for each mouse; p=0.46, paired t-test. Error is S.E.M. across mice. **b)** Same as **a**, for auditory trials; p=0.068, paired t-test. **c)** Reaction time across easy (black) and hard (gray) valid visual and auditory trials; *p<0.05, paired t-test. **d)** Visual trial reaction time across binned target orientation; p<0.01, one-way ANOVA**. e)** Same as **d**, for auditory trials across binned target amplitudes; p=0.66, one-way ANOVA.

